# The Notch1/Delta-like-4 axis is crucial for the initiation and progression of oral squamous cell carcinoma

**DOI:** 10.1101/2024.01.21.576524

**Authors:** Christian T. Meisel, Riccardo Destefani, Ilaria J. Valookkaran, Niels Rupp, Aashil Batavia, Francesca Catto, Jacqueline Hofmann-Lobsiger, Thimios A. Mitsiadis, Cristina Porcheri

**Affiliations:** Institute of Oral Biology, Faculty of Medicine, University of Zurich, Plattenstrasse 11, 8032 Zurich, Switzerland; Department of Pathology and Molecular Pathology, University Hospital Zurich, Rämistrasse 100, 8091 Zurich, Switzerland; IMAI-MedTech, Bio-technopark, Wagistrasse 18, 8952 Schlieren, Switzerland

**Keywords:** Notch signaling pathway, oral cancers, squamous cell carcinoma, OSCC, keratinocyte differentiation, Notch inhibitors, Notch ligands, Delta-like-4, Notch1, Jagged1

## Abstract

The Notch signaling pathway is frequently altered in Oral Squamous Cell Carcinoma (OSCC), the most common malignant neoplasm of the oral mucosa. In this study, we aimed to elucidate the functional role of this pathway in OSCC initiation and progression. Using transgenic animal models, advanced imaging, and next-generation-sequencing techniques, we analyzed Notch-dependent changes driving OSCC. We found specific expression patterns of Notch1 and Delta-like-4 confined to the malignant tissue, while Jagged1 was downregulated in OSCC. This mutually exclusive expression of Delta-like-4 and Jagged1 occurs at the early hyperplastic stage and persists until more advanced stages of the tumor. Transcriptomic analyses confirmed the dysregulation of the Notch pathway circuitry associated with a genetic signature for undifferentiation in OSCC. Furthermore, pharmacological Notch inhibition significantly impaired cancer cell motility. Taken together, these results reveal the pivotal importance of the Notch1/Delta-like-4 signaling axis as a central oncogenic driver in OSCC.

## Main

Head and neck squamous cell carcinoma (HNSCC) is the sixth most common type of cancer, annually affecting 850.000 people worldwide ^1–3^. Although modern multimodal therapy has improved clinical management of this disease, tumor recurrence and distal metastasis correlate with a poor prognosis, with a median overall survival of ten months ^4,5^. HNSCC regroups a variety of cancers arising from the oral mucosa, nasal cavities, pharynx, and larynx, which have their own unique carcinogenic evolution and clinical score ^6^.

In normal mucosal tissues, such as the oral and respiratory epithelia, keratinocytes undergo a well-defined differentiation process as they move from the basal layer to the surface. In the basal layer, stem-cell-like cells express undifferentiated markers such as SRY-box transcription factor 2 (SOX2), P63, and Keratin 5 and 14 (K5, K14). As these cells progress upward they begin to differentiate, producing keratins and junctional and structural proteins essential for the formation of the mucosal barrier ^7^. In SCC, the normal keratinocyte differentiation process is disrupted, leading to uncontrolled proliferation and impaired differentiation ^8^. Basal keratinocytes bypass typical differentiation stages and continue to proliferate in an undifferentiated or partially differentiated state. This results in the accumulation of abnormal, poorly differentiated cells that resist cell death and shedding.

Within tumors, cancer-initiating cells retain self-renewal capabilities, driving tumor initiation and exhibiting resistance to conventional therapies ^9^. These cells share undifferentiated markers with epithelial stem cells, while the expression of keratins indicative of terminally differentiated squamous cells (e.g., K10) is reduced or altered, reflecting the dedifferentiation character of tumor cells.

Oral Squamous Cell Carcinoma (OSCC) originates from the mucosa of the oral cavity, with the tongue being the most commonly affected site. OSCC develops progressively, starting with local hyperplasia of squamous epithelial cells and advancing through dysplastic and invasive stages ^10,3^. In squamous dysplasia, clusters of basaloid nonkeratinizing and poorly differentiated cells grow on the epithelial surface as exophytic growth (e.i. Carcinoma in Situ) ^11^. Aberrant epithelial cells may infiltrate the neighboring muscles and spread along the mesenchymal lines to metastasize to cervical lymph nodes ^4,8^. Proximal metastases occur in up to 50% of patients and are associated with the recurrence and development of distant metastases ^6,12,13^. OSCC onset often relates to chronic exposure to nitrosamines, polycyclic aromatic hydrocarbons, and other genotoxic compounds (e.g. quinoline derivatives, arylamines), which in turn leads to genetic instability and the dysregulation of the DNA repair process^10^. Clones with acquired genetic alterations have an initial growth advantage and can potentially transform into aggressive subclones in the more advanced stages of the disease ^14,15,16^. High-risk premalignant lesions can be subtle, and timely recognition relies on the detection of stage-specific molecular biomarkers.

Mutational profiling consistently identified chromosomal alterations resulting in the amplification of *cyclin D1* and EGFR regions, while genetic mutations affecting tongue epithelium have been mapped in cyclin-dependent kinase inhibitor 2A (CDKN2A), tumor protein 53 (TP53) ^3,17,18^. Notably, the Notch signaling pathway is one of the major mutation hotspots in OSCC^19–21^. Exome sequencing screenings revealed that approximately 15% of tumors harbor mutations in this pathway, including truncations, insertions-deletions, and missense variants.

Notch signaling is an evolutionary conserved pathway involved in cell fate specification and differentiation ^22^. It plays a crucial role in maintaining epithelial integrity, particularly in the oral cavity, where it regulates mucosal formation and homeostasis from embryonic development onward ^22–27^. In mammals, four Notch receptors (Notch1-4) and five Notch-ligands (Jagged1,2 and Delta-like-1,3,4) have been identified ^28,29^. Upon receptor-ligand binding, a series of cleavages take place to ultimately release the Notch intracellular domain (NICD), which then translocates into the nucleus and acts as a transcription factor in a complex with CSL (also known as RBPj) ^22,30^. The activity of Notch is strictly context-dependent, therefore spatial distribution of each receptor, ligand and downstream targets reflects in a distinct modulation of the pathway, hence influencing tissue homeostasis ^31^. Dysregulation of Notch signaling is found in several human pathologies, and novel therapeutic approaches have been developed in recent years to specifically target Notch components ^31,32^. Although frequently mutated in OSCC^19–21^, the precise role of Notch in the initiation and development of this malignancy remains largely unknown, with contrasting results that are supportive of either an oncogenic or a tumor-suppressive role ^33–37^.

Using *ad hoc* murine models, we have explored the functional role of Notch signaling in the progression of OSCC, from initiation to the most advanced aggressive stages. Our findings reveal a mutually exclusive expression of Dll4 and Jag1 that is consistently maintained across all stages of tumor growth, from precancerous lesions to advanced invasive forms. Transcriptomic analysis further confirmed the downregulation of Notch-specific target genes, alongside dysregulation of genes involved in keratinocyte differentiation and cell-to-cell and cell-to-extracellular matrix (ECM) communication. In parallel, pharmacological modulation of Notch-specific transcription in human cancer cells led to a reduction in the malignant phenotype. Together, our data underscore the critical role of the Notch1/Dll4 signaling axis in OSCC initiation and progression and substantiate its suitability as both an early diagnostic tool and a functional target for OSCC treatment.

## Materials and Methods

### Animals

C57Bl/6 and *Notch1^CreERT^:R26mTmG^fl/fl^*mice were used for this study ^38,39^,The mice were maintained and handled according to the Swiss Animal Welfare Law. This study was approved by the Cantonal Veterinary office, Zurich (License ZH197/2017 and ZH086/2021).

### 4NQO treatment of mice – OSCC murine model

For the induction of OSCC, treated mice received drinking water containing 50μg/mL 4NQO (#N8141, Sigma-Aldrich) starting at week 6 until week 22 of age. Control mice were littermates receiving drinking water containing the same volume of DMSO (#D4540, Sigma-Aldrich). The drinking solution was renewed twice a week. Females and males were equally represented in the experimental set-up. After 16 weeks mice were either euthanized (time point 22 weeks) or given normal drinking water until euthanasia (time point 32 weeks or 38 weeks). For lineage tracing Notch1-driven expression, Cre was activated by the intraperitoneal injection of tamoxifen 50mg/kg dissolved in corn oil/10% EtOH (Sigma-Aldrich Cat#T5648) for two consecutive days. All analyses were conducted in groups of 3 to 5 mice (n ≥ 3).

### Sample embedding and cryo-sectioning

After tissue isolation, samples were post fixed in 4% Paraformaldehyde (PFA) for 1h at 4°C and cryopreserved in 30% sucrose. Tissues were embedded in a cryo-embedding matrix (Tissue-Tek, O.C.T Compound, Sakura) and stored at -80°C until sectioning. Samples were sectioned (10μm) in the cryostat (Leica CM3050S) and stored at -80°C until staining.

### Hematoxylin & Eosin staining

Sections were washed for 3x5min in 1x PBS. Nuclei were stained with Hematoxylin for 20 seconds (Sigma-Aldrich #51275). Cytoplasm was then stained with Eosin for 10 seconds (Sigma-Aldrich #HT110216), thoroughly washed and mounted for imaging. Images were acquired using the Leica DMC5400 microscope.

### Immunostaining of cryosections

Immunofluorescence staining of cryosections was performed as previously described^40^. Briefly, Cryosections (10–25 µm thick) were post-fixed with 4% PFA and permeabilized with a 0.5% Triton/PBS solution for 30 minutes at room temperature. To minimize nonspecific antibody binding, slides were then incubated for 1 hour at RT in a blocking buffer containing species-specific serum and 0.1% Triton in PBS. Primary antibodies were diluted in the blocking buffer at the following concentrations: polyclonal rabbit anti-Keratin14 (1:100; Poly19053, Biolegend), polyclonal goat anti-E-Cadherin (1:200, AF748, R&D Systems), polyclonal goat anti-GFP (1:100; ab6673; Abcam), polyclonal rabbit anti-DLL4 (1:100; ab7280; Abcam), polyclonal goat anti-DLL4 (1:50; AF1389; R&D) polyclonal rabbit anti-Jag1 (1:100; ab7771; Abcam), polyclonal rabbit anti-Keratin10 (1:100; PRB-159P; Biolegend), polyclonal Rabbit anti-Keratin5 (dilution 1:100; ab64081; Abcam) polyclonal rabbit anti-Notch1 (D1E11) (dilution 1:100; 3608S; Cell Signaling), polyclonal rabbit anti-Runx1 (dilution 1:100; HPA004176; Atlas Antibodies), polyclonal rabbit anti-p63 (dilution 1:100; ab53039; Abcam), monoclonal rabbit anti-Vimentin (dilution 1:100; ab92547; Abcam), mouse anti-Snail (1:100, #14-9859-80 Invitrogen). Used secondary antibodies: Goat anti-Rabbit IgG 488, Alexa Fluor Plus 488 (dilution 1:1000; A32731, Invitrogen), goat anti-mouse IgG 647, Alexa Fluor Plus 647 (dilution 1:1000; A32728, Invitrogen), donkey anti-Goat IgG 488, Alexa Fluor Plus 488 (dilution 1:1000; A32814; Invitrogen); donkey anti-Rabbit IgG 647, Alexa Fluor 647 (dilution 1:1000; A32795: Invitrogen), streptavidin Alexa Fluor 488 (dilution 1:100; S11223; Invitrogen). Nuclear counterstaining was performed using DAPI (4’,6-Diamidino-2-Phenylindole, Dihydrochloride, #D1306, Invitrogen).

### Image acquisition – Confocal microscopy

Immunofluorescence images of the tongue tissue were produced using a Leica DM6000B microscope, or Confocal Leica SP5/ SP8 with the following settings: 1024x1024 pixel dimension; 400Hz laser; frame average 3 or 4. photomultipliers PMT or Hyd. Fluorochromes used: Alexa 488, 546, 594, 647. A range of 10μm (standard) to 70μm (whole mount) thick Z-stack was analyzed collecting images every 2μm (standard) or 10μm (whole mount). Images were processed using Imaris 9.9 software (Bitplane). Photomerge Script from Photoshop (Adobe, Version 23.0.2) was used for overview images reconstruction.

### Tissue clearance

The separated epithelial sheet was embedded in 2% low-melting agarose. The tissue was then transferred into Histodenz (#D2158, Sigma-Aldrich), dissolved in PBS1X and diluted until a measurable refractive index of 1.457 for tissue clearance.

### Whole-organ 3D imaging

Whole tongue images were recorded with a custom-made selective plane illumination microscope (mesoSPIM) ^41^. SPIM imaging was done after clearing and refractive index matching. The laser/filter combinations for mesoSPIM imaging were as follows: for Notch1-GFP at 488 nm excitation, a 405-488-561-640 nm QuadrupleBlock bandpass filter was used as the emission filter; for membrane localized dt-Tomato at 561 nm excitation, a 561 nm long pass filter was used. Transparent tongue epithelial whole-mount was imaged at a voxel size of 3.26 × 3.26 × 3 µm (X × Y × Z). Further technical details of the mesoSPIM have been previously reported ^41^ (mesoSPIM.org).

### Enzymatic separation of keratinized epithelium from mesenchyme of the murine tongue

Separation was performed as previously described ^42^. In brief, an enzymatic cocktail of collagenase A (1mg/mL; Roche) and Dispase II (2mg/mL; #D4693 Sigma-Aldrich;) in 0.1 M PBS1X was injected into the subepithelial space from the posterior cutting end of the tongue. Upon incubation at 37°C for 30 min the mesenchyme and epithelium were pulled in opposing directions until complete separation is achieved.

### Oral mucosal organoid establishment and treatment

Oral mucosal organoids were established and propagated as previously described ^43^. Briefly, following enzymatic separation of the tongue epithelium, a keratinocyte cell suspension was obtained and cultured in Advanced DMEM^+++^ supplemented with B27 (Life Technologies, #17504-044), 1.25 mmol/L N-acetyl-L-cysteine and 10 mmol/L nicotinamide. Every three days freshly prepared mRSPO1 (100 ng/mL, PeproTech), mEGF (50 ng/mL,), and 10 ng/mL FGF10 was added. Organoids were passaged every 8 days by dissociating them into single cells using a combination of trypsin digestion and mechanical disruption via pipetting. Trypsin was inactivated using 10% FBS for 10 minutes at 37 °C, after which cells were centrifuged and resuspended in complete organoid medium with growth factors. For treatment, either CB103 (100 μM; MedChemExpress, #HY-135145) or an equivalent volume of DMSO (Sigma-Aldrich, #D8418) was added to the organoid medium. Treated single cells were imaged every day, and organoid size was quantified using FIJI/ImageJ software. Quantitative analyses were performed on 40 organoids per condition.

### RNA isolation epithelial tissue

Separated epithelial sheets were snap-frozen with liquid nitrogen and homogenized on ice for 3min using a motor driven grinder (Pellet Pestle Motor, Kontes). Further isolation according to manufacturer’s instructions (RNeasy Mini Kit; Qiagen). Quantification of the total RNA extracted by NanoDrop (Thermo Fischer Scientific, Waltham, USA). RNA was retrotranscribed using the iScript™cDNA synthesis kit (Bio-Rad Laboratories AG, Cressier FR, Switzerland) according to manufacturer’s instructions.

### Quantitative real-time PCR

The quantitative real-time PCR was performed using the CFX Connect Real-Time System and the software CFX Manager Version 3.1. Expression level analysis of *Actin* (housekeeping gene), and all candidate genes were performed using the SYBR^®^ Green PCR Master Mix (Applied Biosystems, Carlsbad CA, USA) in combination with corresponding oligonucleotide primers (Supplementary Table 1 and 2).

All amplifications were done in technical duplicates of 5 biological replicates. Using the following thermocycling conditions: 95°C for 10min; 40 cycles of 95°C for 15sec; 60°C for 60sec. Melting curve analysis was performed at 95°C for 15sec, 60°C for 5sec and 95°C for 30sec. Expression levels were calculated using the ΔΔCt method (2^−ΔΔCt^ formula), after being normalized to the Ct-value of the *Actin* housekeeping gene.

### Bioinformatic analysis

Library for bulk seqRNA was prepared using Illumina NovaSeq 6000. Differentially expressed genes were obtained from the DEseq2 output (p.adj <= 0.05) (Supplementary Method 1 and Supplementary Method 2). We determine pathway enrichment using the ReactomePA v1.38.0 package ^44^. We produce a similarity matrix using the Jaccard distance between the enriched terms needed to produce the enrichment map using the R package enrichplot v1.14.1.

Raw data available on NCBI/SRA database (https://www.ncbi.nlm.nih.gov/sra/PRJNA1067290 and https://www.ncbi.nlm.nih.gov/sra/PRJNA1187878).

### Patients’ samples

Samples from patients diagnosed with HPV-negative HNSCC were used under the license BASEC 2018-01729. Tissues from patients 18-90 years old with primary tumor occurrence were collected upon primary surgical therapy performed at the University Hospital Zurich (USZ). Tissues were processed, embedded in OCT and sectioned according to USZ SOPs. For immunohistochemistry staining, primary antibody α-Jagged1 (1:100; Abcam, ab7771) and α-Dll4 (1:100; R&D Systems, AF1389) were used.

### Cell culture

The SCC-25 epithelial cell line, derived from the tongue of a 70 years-old OSCC patient, was purchased from the ATCC/LGC company (SCC-25CRL-1628™) and maintained accordingly to producer indications. Briefly, cells were cultured in DMEM/F12 medium containing 1.2 g/L sodium bicarbonate, 2.5 mM L-glutamine, 15 mM HEPES, 0.5 mM sodium pyruvate (# 11330032, Gibco-ThermoFisher) and 10% fetal bovine serum (#16000044, Gibco-ThermoFisher). Cells were sub-cultured when 80% confluent using trypsin/EDTA (#15400054, Gibco-ThermoFisher) and frozen at 10^6^ cells/vial in 10% DMSO (#D4540, Sigma-Aldrich). All cells used in our experiments are mycoplasma-free.

### Scratch assay

For scratch assay, SCC25 cells were grown to confluence. A central scratch of 1mm of distance was produced using a silicon scraper. Medium was changed immediately with conditioned medium: Control group medium contained the anti-mitotic Aphidicolin (1ug/ml; # A4487 Sigma-Aldrich) and DMSO (#D8418; Sigma-Aldrich); Treated group contained Aphidicolin (1ug/ml; # A4487 Sigma-Aldrich) and CB103 (100μM; # HY-135145 MedChemExpress). Fresh conditioned medium was used for medium changes every two days for 10 days. Images were taken at 20X every day at the same time of the day using a Leica DMIL LED inverted microscope mounting a Leica camera DFC295. Four wells per conditions were analyzed. Quantification of cell-free areas was performed using FIJI/ImageJ free software.

### Cell immunofluorescence

Treated SCC25 cells were plated on hatched coverslips at 20.000cells/well. Cells were fixed with PFA4% for 15min at 4°C. After PBS washing, cells were permeabilized 30 min in 0.1% triton and then blocked using a solution of 10% FBS, 0,1% triton in PBS. Primary antibodies were diluted in blocking buffer and incubated overnight. Primary antibodies a-Vinculin 1:500 (#90227 Millipore) was used for immunofluorescence on cells and detected via secondary antibody donkey anti-mouse IgG 546, Alexa Fluor 546 (dilution 1:1000; Invitrogen). Nuclear counterstaining was performed using DAPI (4’,6-Diamidino-2-Phenylindole, Dihydrochloride, #D1306, Invitrogen).

TRITC-conjugated Phalloidin staining of actin filaments was performed as indicated by the manufacturer (1:1000 for 1h incubation at room temperature; Millipore # 90228). Mounting in Mowiol 4-88/Glycerol (#81381 and #G-6279, Sigma-Aldrich).

### Time-lapse imaging

Low passage SCC25 cells were treated with CB103 (100μM; # HY-135145 MedChemExpress) or DMSO (#D8418; Sigma-Aldrich) for 4 days and then plated at 10000 cells/well in an 8-wells μ-Slide high glass bottom (#80807, Ibidi). Cells were observed under an Olympus ScanR HCS at 20X with controlled environmental conditions (CO2 5% and 37°C) for 3h, taking an image every 3 minutes. For analysis, Imaris 9.9 (Bitplane) spot and tracking function was used to automatically quantify the displacement length.

## Results

### Physiological Distribution and Localization of Notch1, Dll4, and Jag1 in the Tongue Epithelium

To investigate how the Notch signalling pathway is activated in OSCC, we initially sought to examine receptor-ligand interactions in the keratinized epithelium of the tongue, the primary site of OSCC development. Since different receptor-ligand combinations can either activate or inhibit Notch signaling, identifying the specific interactions present in normal tissue is a critical first step in understanding how the pathway is regulated in the oral mucosa.

To map Notch1 receptor activity in vivo, we used an inducible transgenic reporter mouse model, *Notch1^CreERT^:R26mTmG^fl/fl^*, which allows for mGFP labeling of Notch1-expressing cells following tamoxifen administration (Figure 1A). This system enabled us to monitor Notch1 activation over time and to track the behavior and contribution of labeled cells as the tissue transitions from a healthy state to advanced stages of OSCC.

**Figure 1:**
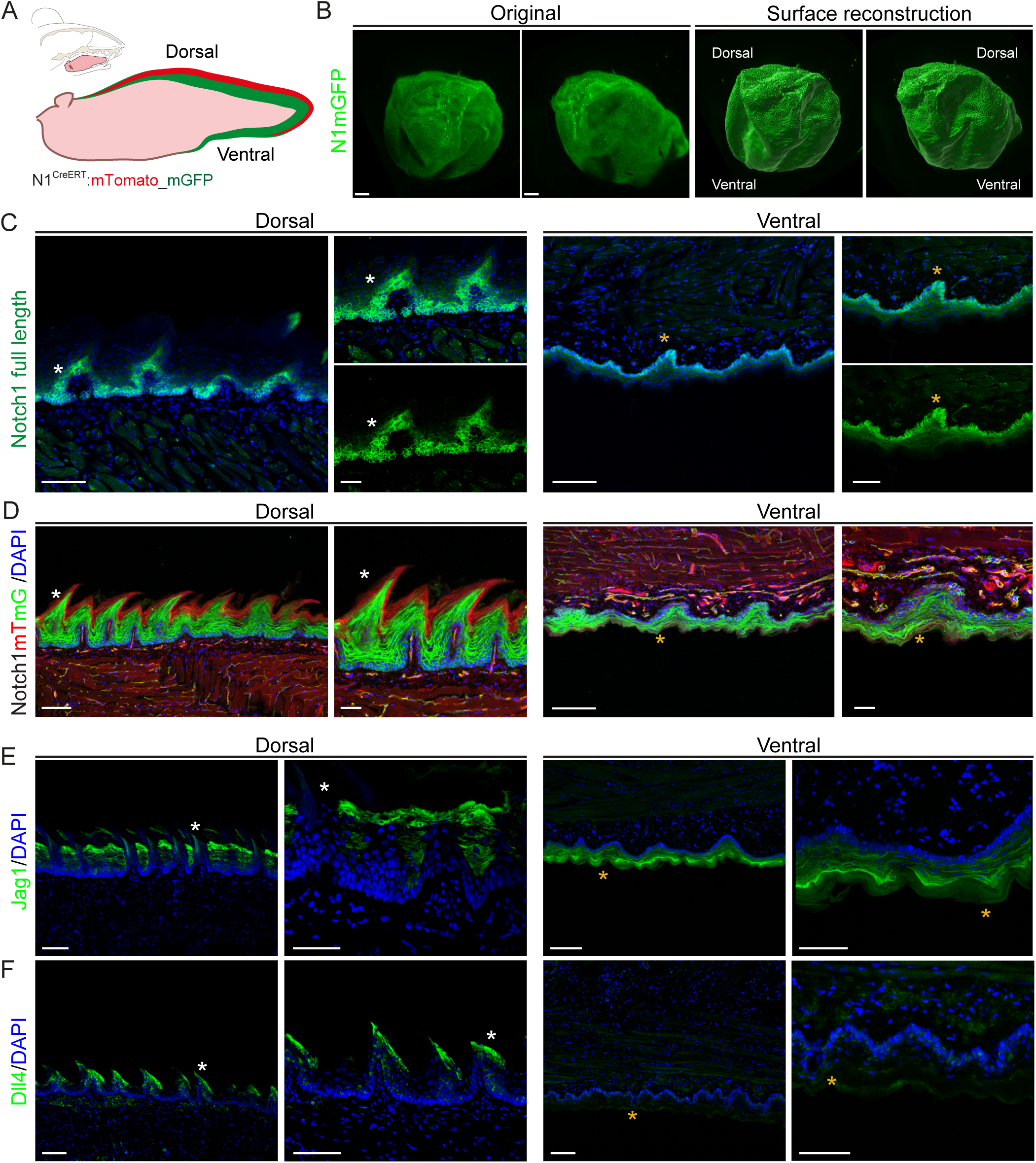
Notch1, Dll4 and Jag1 expression in the epithelium of the mouse tongue. A) Schematic representation illustrating the murine tongue of a *Notch1^CreERT^:R26mTmG^fl/fl^* mouse after tamoxifen induction and subsequent *mGFP* expression. B) MesoSPIM imaging and related three-dimensional reconstruction of a whole-mount *Notch1^CreERT^:R26mTmG^fl/fl^*mouse tongue. Notch1-driven *mGFP* expression (green) in the raw reconstruction (left panels) and relative surface rendering (right panels). C) Cross-sections of the dorsal (left panels) and ventral (right panels) part of the tongue stained against the full length Notch1 protein (green). Cell nuclei are stained with DAPI (blue). Asterisks indicate the magnified areas in the corresponding region of the overview. D) Cross-sections showing the dorsal (left panel) and ventral (right panel) parts of the tongue of a *Notch1^CreERT^:R26mTmG^fl/fl^*mouse. Both the dorsal and the ventral epithelium express the Notch1-driven mGFP (green). Unrecombined tissue expresses *mTomato* (red). Cell nuclei stained with DAPI (blue). Asterisks indicates magnified area in the corresponding region of the overview. E) Cross-sections of the dorsal (left panel) and ventral (right panel) parts of the tongue showing Jag1 staining (green). Cell nuclei are stained with DAPI (blue). F) Cross-sections of the dorsal (left panel) and ventral (right panel) parts of the tongue showing Dll4 staining (green). Nuclei are stained with DAPI (blue). Scale bars in overviews100μm. Scale bars in magnification 50μm.

Using light-sheet microscopy (mesoSPIM), we generated a 3D reconstruction of the entire tongue epithelium and observed Notch1-driven mGFP expression in both ventral and dorsal regions (Figure 1B, Movie 1). Cross-sections of adult tongues provided more detailed views of Notch1 localization within the stratified epithelium, confirming its broad distribution across both surfaces (Figure 1C). In *Notch1^CreERT^:R26mTmG^fl/fl^*mice, Notch1:mGFP-positive cells were restricted to the multilayered tongue epithelium, while unrecombined regions retained expression of the mTomato cassette (Figure 1D). Both endogenous Notch1 protein and Notch1-driven mGFP were detected from the basal layer up to the stratum granulosum (Figures 1C, 1D).

We then examine the spatial distribution of Notch-ligands Jagged1 (Jag1) and Delta-like-4 (Dll4) in the tongue epithelium. We identified a compartmentalized expression pattern with mutually exclusive localization (Figures 1E and 1F). Jag1 was present on both the dorsal and ventral sides of the tongue, with staining in the spaces between filiform papillae of the dorsal region (Figure 1E). In contrast, Dll4 was exclusively localized in the epithelium of the dorsal tongue, specifically on the outer surface of the filiform papillae (Figure 1F). This complementary expression pattern of the two ligands suggests a potentially distinct role for Notch1-Jagged1 versus Notch1-Dll4 interactions in the physiology of tongue keratinocytes.

### Mutually exclusive distribution patterns of Dll4 and Jag1 in the tongue epithelium of OSCC

To model the progressive stages of tongue squamous cell carcinoma, we administered the carcinogenic substance 4-Nitroquinoline N-oxide (4NQO) in the drinking water of 6-weeks-old mice for a duration of 16 weeks (Supplementary Figure 1A) ^45^. As previously reported, chronic exposure to 4NQO for 16 weeks induces hyperplasia and dysplasia in the tongue epithelium ^46^. After an additional 10-week washout period, the tissue progresses to carcinoma with 100% penetrance. Histological analysis of 22-week-old mice revealed epithelial hyperplasia, leukoplakia, and hyperkeratinization, primarily on the dorsal side of the tongue and occasionally on the ventral side (Supplementary Figures 1A, 1B, 1C). By 32 weeks of age, histological evidence of carcinoma in situ was observed on the dorsal epithelium of the tongue (Supplementary Figures 1B, 1C) ^46^

To study the role of Notch in OSCC initiation, we firstly treated *Notch1^CreERT^:R26mTmG^fl/fl^* mice with 4NQO or DMSO and analyzed the tissue at 22 weeks of age (Figures 2A). In 4NQO-treated mice, we observed specific Notch-driven mGFP expression localized to the hyperplastic epithelium (Figure 2B), indicating active Notch signaling within the cell population driving malignant transformation. Next, we assessed the distribution of the Notch ligands Dll4 and Jag1 at this early stage. Dll4 staining was restricted to the hyperplastic epithelium (Figure 2C), while Jag1 immunolabelling was notably absent in this same region (Figure 2D, white arrows and dashed area). Interestingly, Jag1 expression displayed a gradual decline: it remained at physiological levels in unaffected tongue regions distant from the epithelial overgrowth (Figure 2D, red arrows), decreased progressively in the tissue bordering the hyperplastic area (Figure 2D, yellow arrows), and became undetectable within the tumor’s central core (Figure 2D, white arrows).

**Figure 2:**
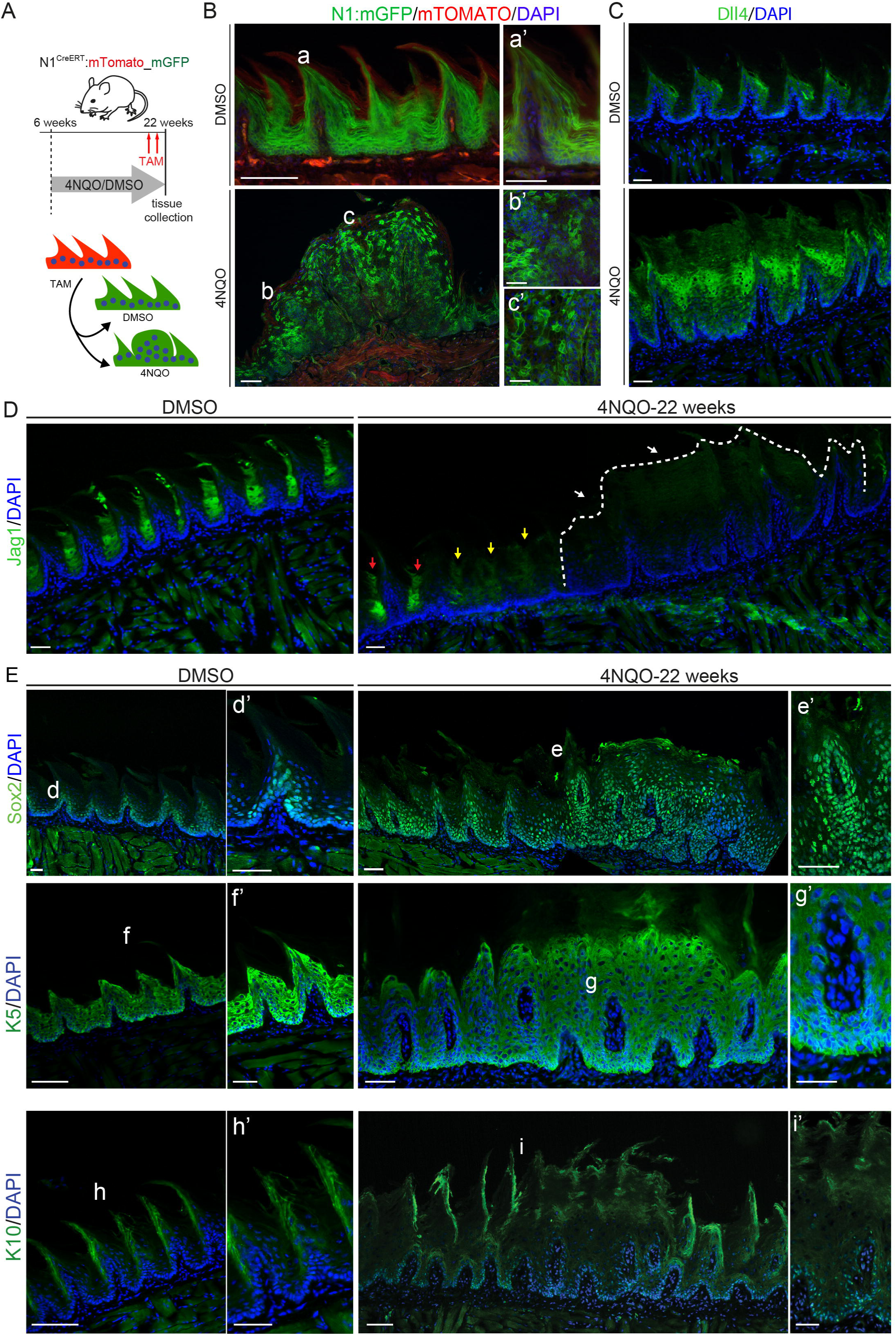
The Notch ligand Jagged1 and Dll4 expression is mutually exclusive in the early stage of OSCC. (A) Schematic representation of the tamoxifen induction and the 4NQO or DMSO treatment in 22-week-old *Notch1CreERT:R26mTmGfl/fl* mice. (B) Cross-sections of the dorsal part of the tongue in DMSO-treated (upper panels) and 4NQO-treated (lower panels) mice. mTomato expression (red) and mGFP expression (green) in the healthy (DMSO) and aberrant (4NQO) epithelium of the tongue. Cell nuclei are stained with DAPI (blue). (a-c) indicate the magnified areas in the corresponding regions of the overview (a-c’). (C) Cross-sections of the dorsal part of the tongue stained against Dll4 (green) in DMSO- (upper panels) and 4NQO-treated (lower panels) mice. Cell nuclei are stained with DAPI (blue). (D) Cross-sections of the dorsal part of the tongue stained against Jag1 (green) in DMSO-treated (left panels) and 4NQO-treated (right panels) mice. Cell nuclei are stained with DAPI (blue). Note the gradient of the Jag1 staining in the 4NQO-treated epithelium: the epithelium distant from the lesion is Jag1-positive (red arrows), the staining is reduced in the epithelium close to the hyperplasia (yellow arrows) and is absent in the hyperplastic tissue (white arrows). The dotted line indicates the borders of the epithelial hyperplasia. (E) Cross-sections of the dorsal part of the tongue showing Sox2 staining (green, upper panels), K5 (green, middle panels) and K10 (green, lower panels) in the healthy (DMSO) and aberrant (4NQO) epithelium of the tongue. Cell nuclei are stained with DAPI (blue). (d-i) indicate the magnified areas in the corresponding regions of the overview (d’-i’). Scale bars in overviews100μm. Scale bars in magnification 50μm.

To determine whether the exophytic growth exhibited undifferentiated characteristics at this early stage of the disease, we performed labeling for the undifferentiated markers Sox2 and K5, alongside the mature keratinocyte marker K10 ^47,48^. Sox2 and K5 were highly expressed in the exophytic mass, whereas K10 was absent, indicating that the growing mass predominantly consists of undifferentiated progenitor cells (Figure 2E).

We next aimed to determine whether the Notch1-Dll4 signaling axis remains active throughout OSCC progression or is restricted to the tumor initiation phase. To address this, we analyzed 4NQO-treated *Notch1CreERT:R26mTmGfl/fl* mice at a later time point (32-weeks-old; Figure 3A). While DMSO-treated mice showed a similar pattern to the one observed in 22-weeks-old tissue (Figure 3B, upper panel), carcinoma in situ lesions exhibited widespread Notch1-driven mGFP labeling (Figure 3B, lower panel). Consistent with our previous observations, Jag1 expression was largely absent from the tumor mass but persisted in the adjacent tissue (Figures 3C), whereas strong Dll4 staining was detected within the exophytic tumor growth (Figure 3D). Thus, the Notch1-Dll4 interaction appears to be a constant throughout the course of OSCC development, from initiation to advanced lesions.

**Figure 3:**
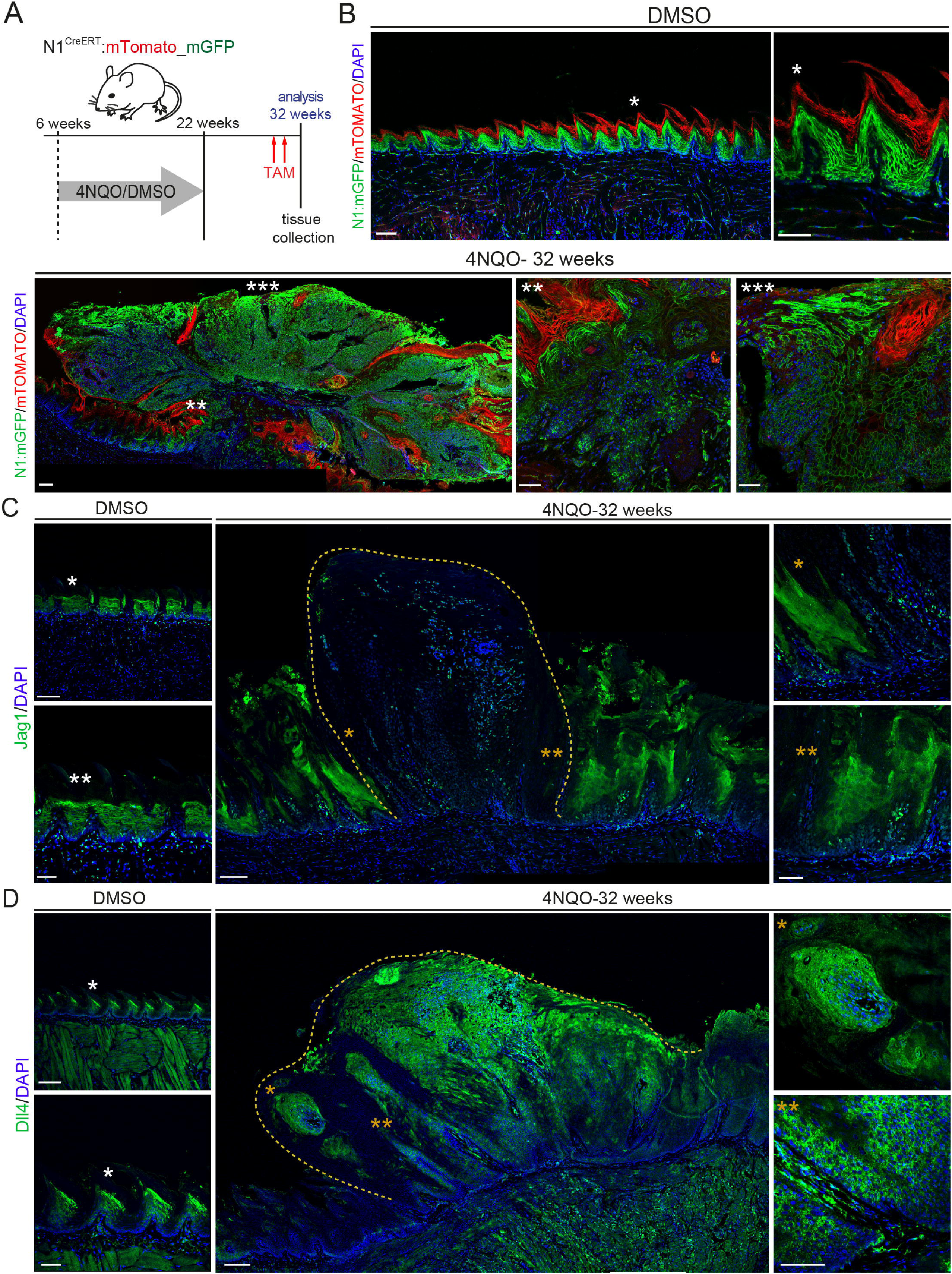
Dll4 is specifically expressed in the OSCC exophytic outgrowth. (A) Schematic representation of the 4NQO or DMSO treatment in 32-week-old *R26:mTomatofl/fl* mice. B) Cross-sections of the dorsal part of the tongue in DMSO-treated (upper panels) and 4NQO-treated (lower panels) mice. *mTomato* expression (red) and *mGFP* expression (green) in the healthy (DMSO) and aberrant (4NQO) epithelium of the tongue. Cell nuclei are stained with DAPI (blue). Asterisks indicate the magnified areas in the corresponding regions of the overview. (C) Cross-sections of the dorsal part of the tongue stained against Jag1(green) in DMSO- (left panels) and 4NQO-treated (right panels) mice. Cell nuclei are stained with DAPI (blue). (D) Cross-sections of the dorsal part of the tongue stained against Dll4 (green) in DMSO-treated (left panels) and 4NQO-treated (right panels) mice. Cell nuclei are stained with DAPI (blue). The dotted line indicates the borders of the OSCC overgrowth. Asterisks indicate the magnified areas in the corresponding regions of the overview. Scale bars in overviews100μm. Scale bars in magnification 50μm.

### Notch1-Dll4 signature correlate with maintenance of undifferentiation and early epithelial to mesenchymal transition

As the Notch signaling have been implicated in the maintenance of stem cell identity in a variety of tissues^49^, we explored whether the presence of Notch1-Dll4 signaling correlates with undifferentiation in OSCC lesions. To this end, 32-week-old mice tissue was stained for the progenitor markers Sox2 and K5. In 4NQO-treated mice, these markers label the majority of malignant cells, while in DMSO-treated controls, Sox2 and K5 staining was restricted to the basal epithelial layer of the tongue (Supplementary Figures 2A and 2B) ^50,51^. Conversely, the differentiation marker K10 was confined in the cornified layer of the healthy tongue epithelium (Supplementary Figure 2C), while remaining undetectable in the 4NQO-induced exophytic growth. Finally, malignant epithelial cells retain the expression of the pan-epithelial marker E-Cadherin (E-Cad)^52^ (Supplementary Figure 2D). Taken together, these results suggest impaired differentiation and an expansion of progenitor-like cells within the 4NQO-induced tumor (Supplementary Figure 2C).

Next, we investigated whether inhibiting Notch in tongue keratinocytes directly affects stem cell behavior ^67^. To assess this, we performed an organoid formation assay using epithelial stem cells isolated from the tongue. In control cultures treated with DMSO, organoids consistently formed within 5 days, with an average diameter of 122.43 μm ± 36.9. However, in cultures treated with the Notch inhibitor CB103, organoid formation was completely disrupted: Cells remained as single cells or small clusters, with an average diameter of 20.21 μm ± 7.19 (DMSO vs. CB103, p ≤ 0.0001). These results suggest that Notch signaling is essential for maintaining the stem cell functions required for organoids initiation and growth (Figure 4A).

**Figure 4:**
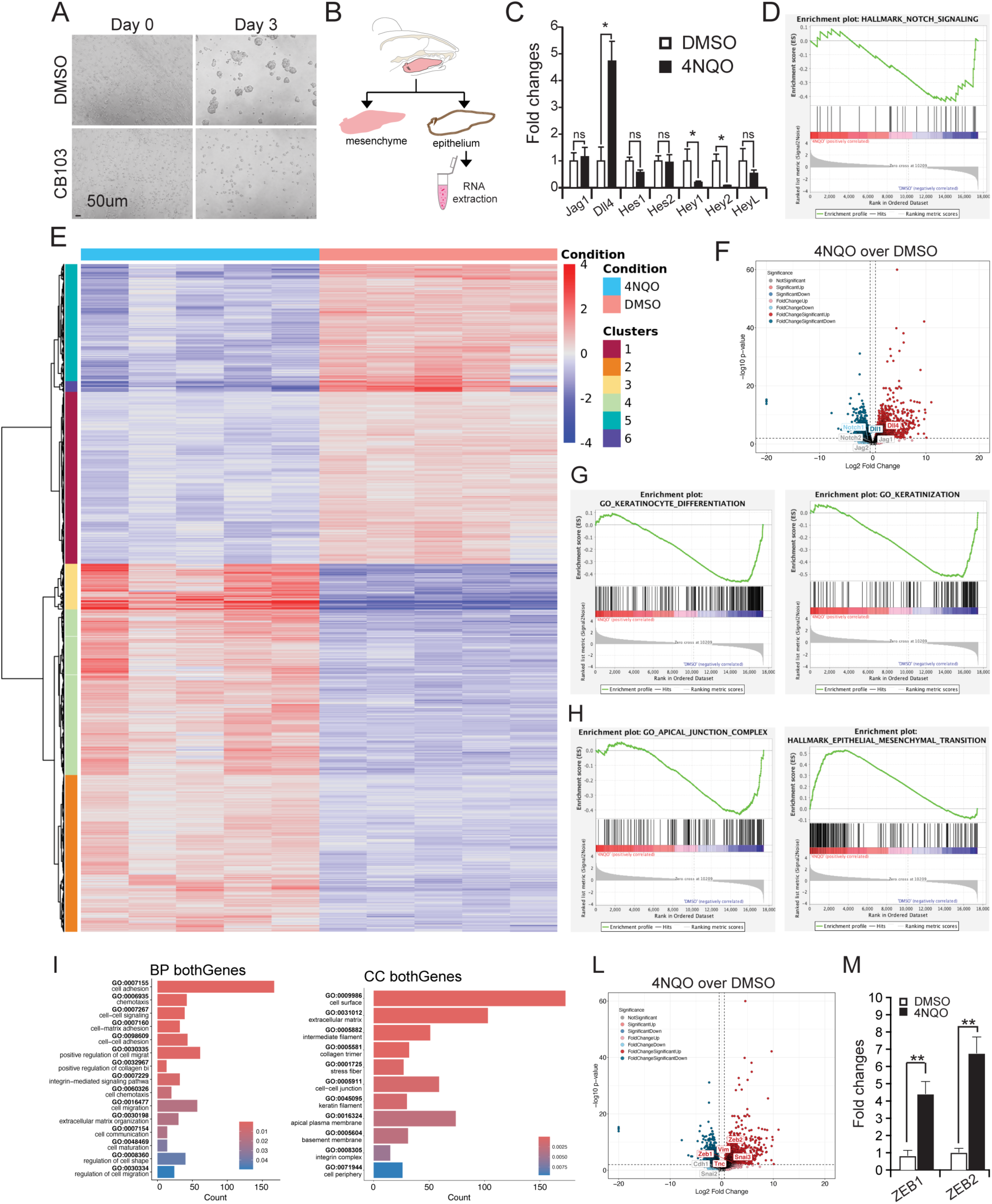
Transcriptome analysis of differential gene expression in the tongue epithelium of 4NQO-treated mice. (A) Organoid formation assay using single epithelial stem cells derived from the tongue. Treatment with CB103 inhibits organoid formation from single-cell suspensions, indicating impaired stem cell function. Scale bar: 50 µm. (B) Schematic representation of the tongue’s epithelium separation for RNA extraction and follow up analysis. (C) qRT-PCR shows upregulation of Dll4 expression and downregulation of Hey1 and Hey2 expression in 4NQO-treated mice. Error bars indicate standard error. *=p<0.05; **=p<0.01; ***=p<0.001 (D) Gene Set Enrichment Analysis (GSEA) plot for the Notch-signaling signature shows changes in the Notch pathway in 4NQO condition when compared to DMSO. (E) Heatmap visualization of 2000 differentially expressed genes with hierarchical clustering. (F) Volcano plot confirm higher expression of Dll4 in 4NQO over DMSO conditions. (G) Gene Set Enrichment Analysis (GSEA) shows alterations in the process of keratinocytes differentiation and keratinization in 4NQO condition. (H) GSEA identify alterations in the apical junction complex and EMT signatures (I) Gene Ontology (GO) classification analysis for categories of biological processes (left panel) and cellular components (right panel) comparing DMSO- versus 4NQO-treated epithelium. (L) Volcano plot identified an increased expression of EMT markers Vimentin, ZEB1, ZEB2, Tenascin C and Snail3. (M) qRT-PCR confirmation of EMT markers ZEB1 and ZEB2 upregulated in 4NQO versus DMSO condition. Error bars indicate standard error. *=p<0.05; **=p<0.01; ***=p<0.001

To gain insight into the molecular changes driving malignant transformation, we performed further analyses at the 32-weeks-old stage. After separating the tongue’s outer epithelium from the underlying connective tissue ^42^ total RNA was extracted for downstream analyses (Figure 4B). qRT-PCR on 4NQO-treated epithelium showed an important increase in *Dll4* expression compared to DMSO-treated epithelium (Figure 4C). Interestingly, the *Hes*-family of target genes remained largely unaffected, whereas *Hey1* and *Hey2* (Hairy and enhancer of split-related with YPRW motif protein) were significantly reduced in 4NQO-treated epithelia (Figure 4C). To obtain a comprehensive view of the transcriptional changes, we additionally performed bulk RNA sequencing analyses. Gene Set Enrichment Analysis (GSEA) revealed a significant enrichment of the Notch signaling signature (Figure 4D), which was further supported by the volcano plot highlighting a pronounced upregulation of *Dll4* in 4NQO-treated samples (Figure 4F). We identified 3331 differentially expressed genes in 4NQO-treated versus DMSO-treated epithelia (p-value threshold p≤ 0.01; Supplementary Method 1). The heatmap shows changes in genes mainly involved in cell differentiation processes (Cluster 2 and Cluster 3), molecular pathways regulating cell fate determination (Cluster 1 and Cluster 5), extracellular matrix rearrangement (Clusters 1,2,3,4), and cell motility (Cluster 3) (Figure 4E, Supplementary Table 3). Additional GSEA showed changes in keratinocyte differentiation and cornification (Figure 4G and Supplementary Figure 3). These results align with our phenotyping data, which show that the Notch1-Dll4 axis is associated with the maintenance of an undifferentiated state.

We then conducted interactome analyses to examine changes in the tongue’s epithelia upon 4NQO and DMSO treatments. Independent *Enrichr (*https://maayanlab.cloud/Enrichr/) analyses identified alteration of *Runt-related transcription factor 1 (Runx1)* amongst others (e.g., *p21, p300* and *Stat3*) (Supplementary Figure 4). Importantly, this alteration is specific to *Runx1*, as other members of the *Runx* family were not significantly altered (Supplementary Figure 4B). As *Runx1* has been previously linked to Notch pathway activation^53–55^, we wanted to demonstrate the distribution of the Runx1 protein in the tumorigenic epithelium of the tongue. Indeed, a strong Runx1 staining was detected in the tumor mass of the 4NQO-treated mice (Supplementary Figure 5A), while the Runx1 labelling was confined to the stratum basale and stratum spinosum of the DMSO-treated mice. Furthermore, strong immunostaining against p63, a marker of undifferentiated keratinocytes^56^, was detected in the tumor mass of 4NQO-treated mice (Supplementary Figure 5B) ^53,57–62^ . Together, these findings suggest that Notch1-Dll4 signaling is closely linked to the expansion of progenitor-like cell populations in OSCC lesions, supporting its role in maintaining an undifferentiated state during malignant transformation.

At more advanced stages of OSCC progression, malignant cells residing within the exophytic mass may become invasive by undergoing epithelial-to-mesenchymal transition (EMT) ^63^. Using light-sheet microscopy (SPIM), we analyzed 4NQO-treated mice to identify the time window when basal lamina dissolution begins. In 22-week-old mice, digital sectioning revealed an uninterrupted basal lamina underneath the epithelial overgrowth (Supplementary Figure 6A, yellow arrows; Movie 2). In 32-week-old mice, we observed a localized area of basal lamina degradation, allowing the malignant overgrowth to directly interface with the underlying mesenchyme (Supplementary Figure 3B, white arrows; Movie 3).

To investigate the molecular changes that may contribute to the dissolution of the basal lamina observed in SPIM imaging, we examined our sequencing data. We identified an EMT signature through GSEA, along with changes in anchoring junctions (Figure 4H). Gene Ontology analysis for biological processes and cellular components further confirmed alterations in cell-cell and cell-to-ECM communication, as well as cell migration (Figure 4I). To validate EMT onset at this stage, we assessed the expression of the mesenchymal marker Vimentin (*VIM*), components of the extracellular matrix Tenascin (*TNC*), and members of the zinc-finger transcription factor family *Snail*. Transcriptomic analyses confirmed their upregulation, together with the early EMT markers ZEB1, ZEB2 (Figure 4L and 4M).

While technical limitations of the 4NQO model prevent us from capturing the full extent of the EMT process and metastasis formation, our data indicate that EMT initiation occurs at the 32-week-old stage following 4NQO exposure, with changes in tissue architecture, molecular signatures, and the upregulation of key markers of cellular transformation.

### Invasive squamous cells conserve their dependency from the Notch1-Dll4 signaling axis

To confirm the progression toward an invasive phenotype, we analyzed a more advanced stage of our OSCC model at 38-weeks-old. Tongue sections from *R26:mTomato* mice were stained with the epithelial markers K14 and P63. This labeling revealed clusters of epithelial cells embedded within the stromal tissue of the tongue (Figures 5B and 5C). These findings led us to hypothesize that carcinoma in situ cells had breached the basal lamina and begun invading the underlying connective tissue. As confirmation that EMT was occurring at this stage, we examined the expression patterns of Snai1 and the adhesion molecule regulator E-cadherin ^64,65^. Snail-positive staining was detected in cells within the invasive tumor mass of 4NQO-treated mice (Figure 5D, white arrows), whereas such labeling was absent in DMSO-treated controls. We then assessed the distribution of Vimentin and E-cadherin, markers of mesenchymal and epithelial identity, respectively ^65,66^. A subset of cells within the tumor mass coexpressed vimentin and E-cadherin (Figure 5E, yellow arrows), thus confirming an ongoing EMT process.

**Figure 5:**
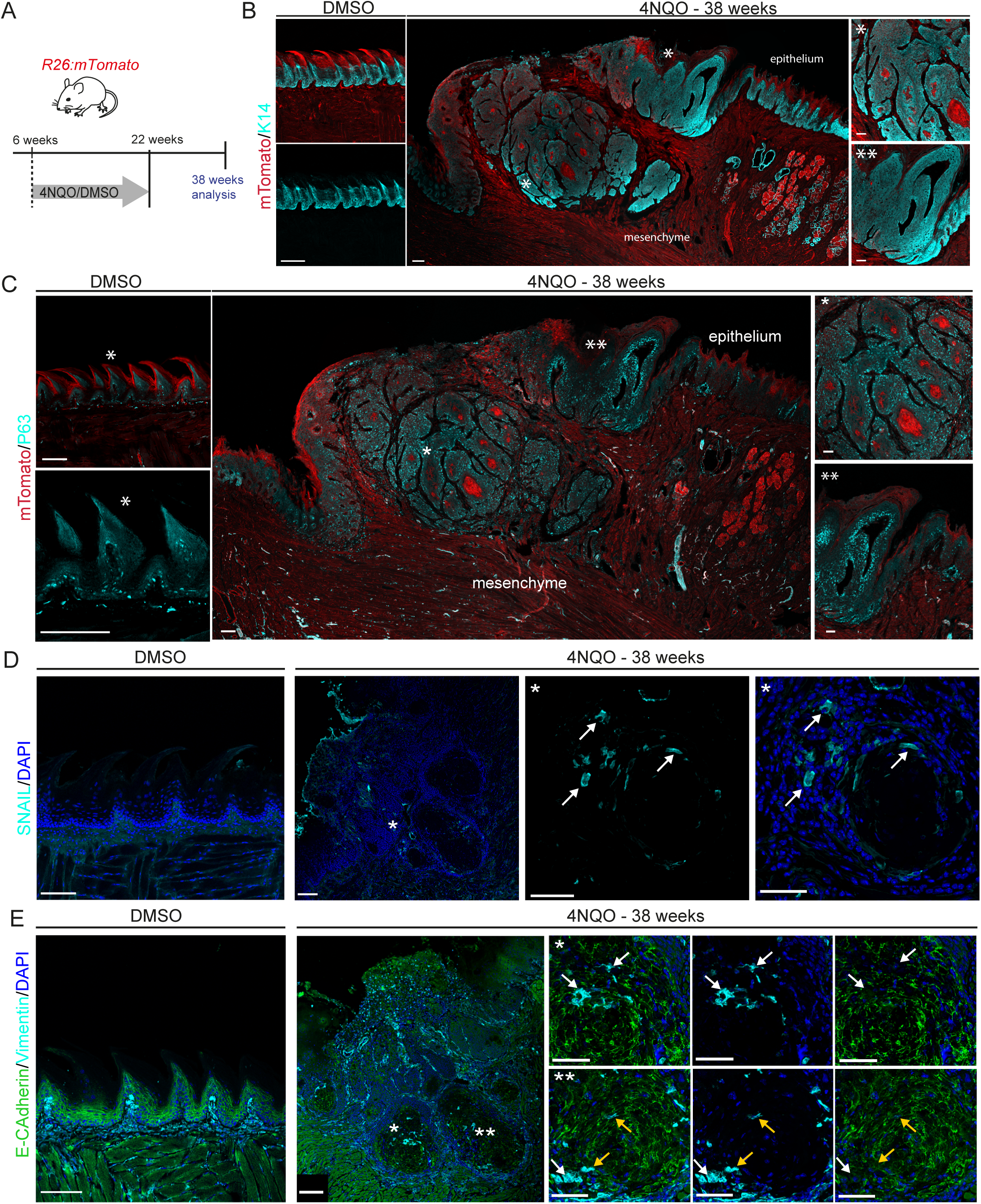
Epithelial cells invade the tongue’s mesenchyme in 4NQO-treated mice. (A) Schematic representation of the 4NQO or DMSO treatment in 38-week-old *R26:mTomatofl/fl* mice. (B) Cross-sections displaying the dorsal part of the tongue labelled with K14 (cyan) in DMSO-treated (left panels) and 4NQO-treated mice (right panels). Background tissue expressed *mTomato* (red). Asterisks indicate the magnified areas in the corresponding regions of the overview. (C) Cross-sections displaying the dorsal part of the tongue labelled with P63 (cyan) in DMSO-treated (left panels) and 4NQO-treated (right panels) mice. Background tissue expressed *mTomato* (red). Asterisks indicate the magnified areas in the corresponding regions of the overview. (D) Cross-sections of the dorsal part of the tongue stained with Snail (cyan) in DMSO-treated (left panel) and 4NQO-treated condition (right panels) mice. Cell nuclei are stained with DAPI (blue). Asterisk indicates the magnified area in the corresponding region of the overview. White arrows indicate Snail-positive cells. (E) Cross-sections of the dorsal part of the tongue stained with Vimentin (cyan) and E-cadherin (green) in DMSO-treated (left panel) and 4NQO-treated (right panels) mice. Cell nuclei are stained with DAPI (blue). Asterisks indicate the magnified areas in the corresponding regions of the overview. White arrows indicate sole Vimentin staining, while yellow arrows indicate staining for both Vimentin and E-cadherin. Scale bars in overviews 100μm. Scale bars in magnification 50μm.

We therefore sought to clarify if the Notch1-Dll4 signaling axis is also involved in the process of tumor invasion observed in 38-weeks-old mice. *Notch1^CreERT^:R26mTmG^fl/fl^* mice were used to trace Notch1-expressing cells and their progeny (Figure 6A). In DMSO-treated mice, *mGFP* expression remained restricted to the epithelial layers (Figure 6B). Conversely, in 4NQO-treated mice, mGFP-labeled cells were observed within the carcinoma mass invading the mesenchyme (Figure 6C). We next performed immunolabeling for the Notch1 protein. A widespread Notch1 signal labeled the infiltrating mass of 4NQO-treated mice, while in DMSO-treated mice Notch1 expression remained confined to the healthy tongue epithelium (Figure 6D). Finally, we examined the expression of Dll4 in the invasive aberrant epithelium using *R26:mTomato* mice. Analysis of DMSO-treated mice confirmed the expression of Dll4 in the dorsal portion of the filiform papillae, as previously observed (Figure 6E and 1F). In 4NQO-treated mice, labeling for Dll4 was broadly present in the infiltrating mass (Figure 6E). Our findings suggest a conserved paradigm governing tumor progression, where the Notch1-Dll4 axis remains central throughout the phases of hyperplastic growth, carcinoma in situ and the most advanced, invasive stages of the disease.

**Figure 6:**
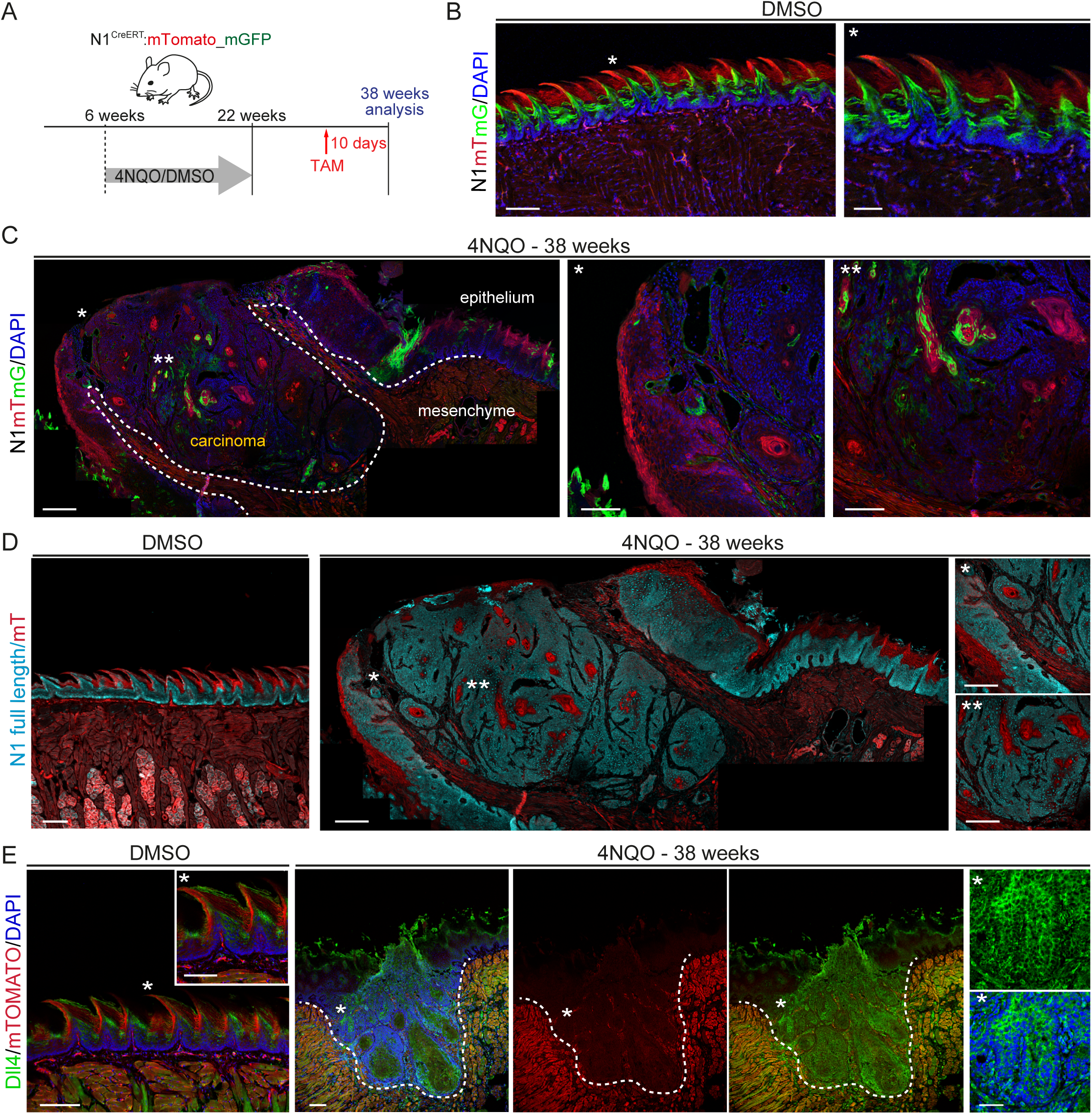
Detection of Notch1-expressing epithelial cells and their progeny within the infiltrated mesenchyme of the tongue. (A) Schematic representation illustrating the tamoxifen induction and 4NQO or DMSO treatment of 38-week-old *Notch1^CreER^T:R26mTmG^fl/fl^* mice. (B) Cross-sections displaying the Notch1-driven expression of *mGFP* in DMSO-treated epithelium. Unrecombined tissue expressed *mTomato* (red). Cell nuclei are stained with DAPI (blue). Asterisk indicates the magnified area in the corresponding region of the overview. (C) Tongue cross-sections displaying the Notch1-driven expression of *mGFP* in 4NQO-treated mice after ten days of tamoxifen induction. Unrecombined tissue expressed *mTomato* (red). Cell nuclei are stained with DAPI (blue). Asterisks indicate the magnified areas in the corresponding regions of the overview. (D) Cross-sections displaying the endogenous Notch1-full length expression (cyan) in the tongue epithelium of DMSO-treated (left panels) and 4NQO-treated (right panels) mice. Unrecombined tissue expressed *mTomato* (red). Asterisks indicate the magnified areas in the corresponding regions of the overview. (E) Cross-sections showing the distribution of Dll4 (green) in the tongue epithelium of DMSO-treated (left panels) and 4NQO-treated (right panels) mice. Asterisk indicates magnified area in the corresponding region of the overview. Dashed lines indicate the extent of the epithelial infiltration in the musculature of the tongue. Scale bars in overviews 100μm. Scale bars in magnification 50μm.

### The Notch signaling pathway regulates malignancy in human OSCC

To evaluate the clinical relevance of the Notch pathway in OSCC, we examined the expression of JAG1 and DLL4 in human tissue samples. We first performed Hematoxylin and Eosin (H&E) staining on tissues from HPV-negative adult patients with primary tumors. This allowed us to identify characteristic SCC features, including dysplastic epithelium, mitotic figures, and abnormal keratin accumulation (Figure 7, overview). Based on these histological criteria, we distinguished healthy tissue (Figure 7A) from hyperplastic (Figure 7D) and carcinoma samples (Figure 7G).

**Figure 7:**
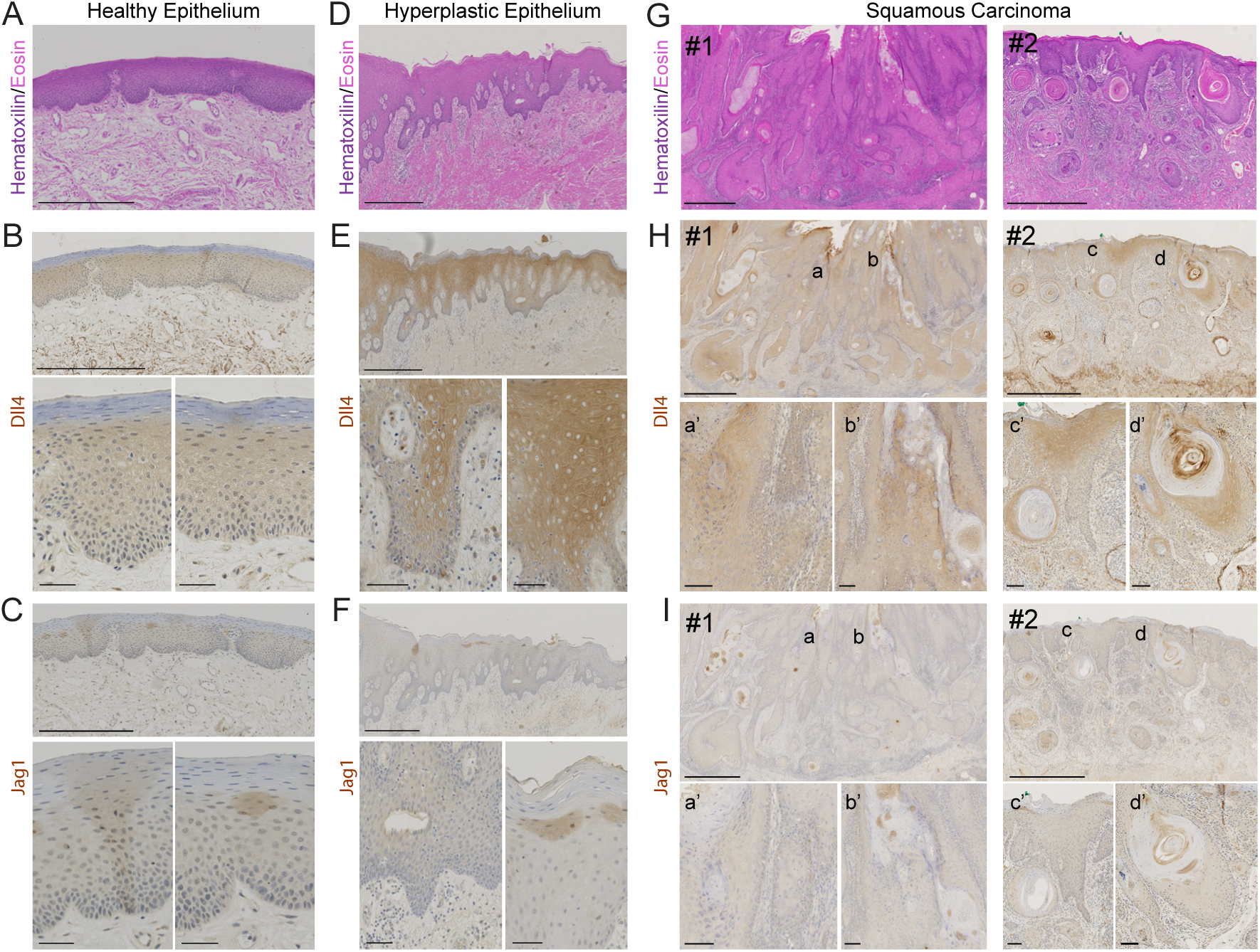
Notch signaling in human squamous cell carcinoma. (A) Histological examination of human healthy epithelium. Hematoxylin/Eosin staining is used to identify the histological condition of the samples analyzed. Corresponding areas are then stained for Dll4 (B) and Jag1(C). Overview of expression for Dll4 (upper panel) and magnifications (lower panels). Scale bar in overview 500μm. Scale bar in magnification 50μm. (D) Histological examination of human hyperplastic epithelium. Hematoxylin/Eosin staining is used to identify the histological condition of the samples analyzed. Corresponding areas are then stained for Dll4 (E) and Jag1(F). Overview of expression (upper panel) and magnifications (lower panels). Scale bar in overview 500μm. Scale bar in magnification 50μm. (G) Histological examination of human squamous cell carcinoma tissue. Hematoxylin/Eosin staining is used to identify aberrant histology in two representative patients (Patient#1 with hyperplastic epithelium: middle panel, Patient#2 with squamous carcinoma: right panel). (H) Distribution of Dll4 in Patient#1 and Patient#2. Magnified areas show expression of Dll4 protein. Scale bar in magnification: 100μm. Scale bar in overview: 1mm. (I) Distribution of Jag1 protein in Patient#1 and Patient#2 in overview and magnified areas. Scale bar in magnification: 100μm. Scale bar in overview: 1mm.

Jag1 and Dll4 expression were then compared across these different tissue types. DLL4 showed diffuse expression in the healthy epithelium (Figure 7B), with increased levels observed in hyperplastic tissue (Figure 7E) and consistently high expression in carcinoma nests (Figure 7H). In contrast, JAG1 staining in healthy tissue was limited to small, localized pockets, and its expression varied across different regions of the carcinoma samples (Figure 7C and Figure 7F,7I).

To further investigate the molecular regulation of the Notch pathway in the human context, we treated hSCC25 cell line (#CRL-1628, ATCC/LGC) with CB103, a pan-blocker of the Notch-transcriptional activation complex^67^ (Figure 8A). Interestingly, quantitative RT-PCR revealed a specific downregulation of the *DLL4* expression, while *JAG1* levels remain largely unchanged (Figure 8B). Additionally, CB103 treatment resulted in decreased expression of *SOX2* and *P63* (Figure 8B), suggesting a direct link between Notch activity and the maintenance of an undifferentiated state.

**Figure 8:**
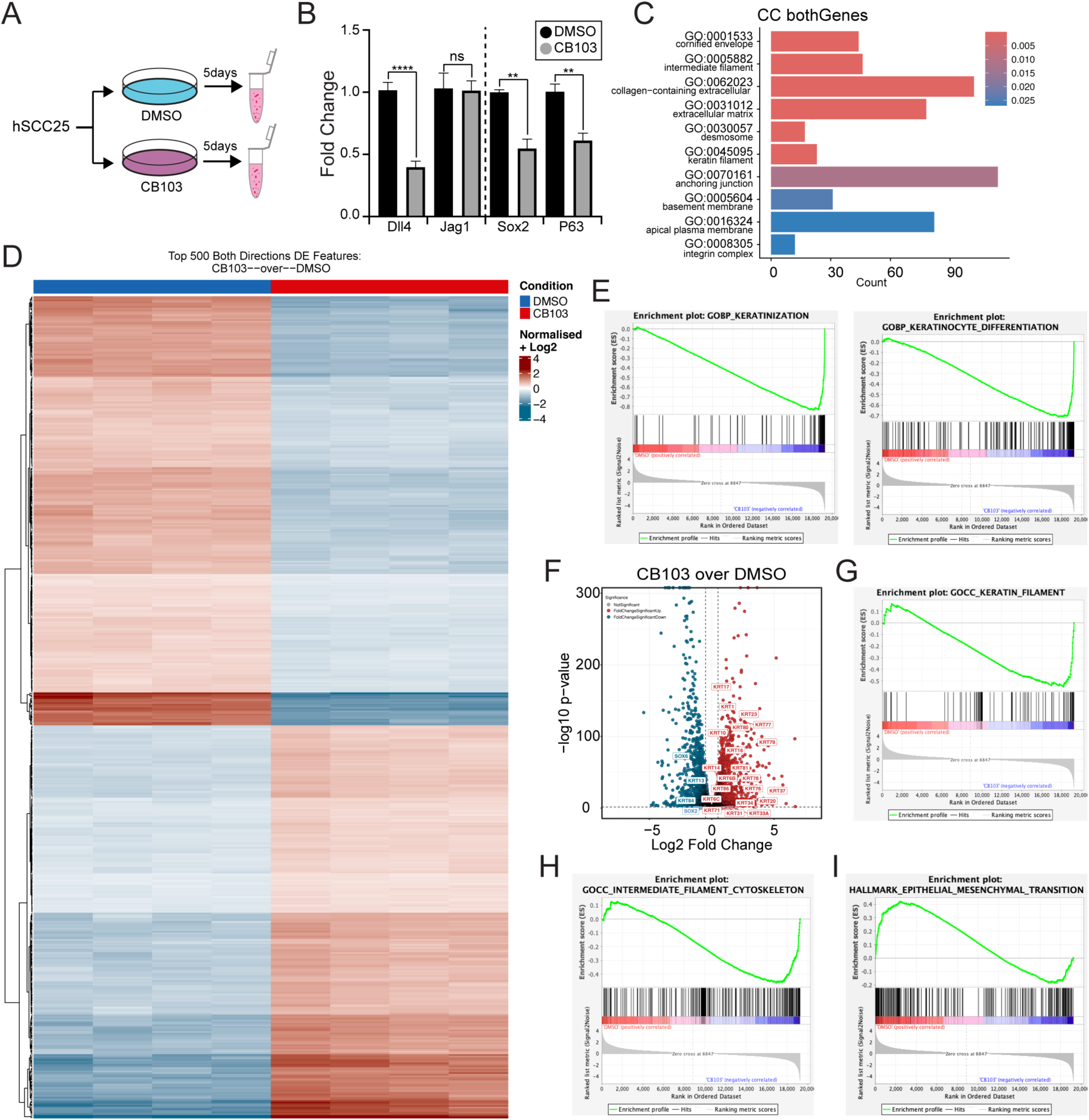
CB103 treatment leads to broad transcriptional reprogramming in human squamous cell carcinoma cells. (A) Schematic overview of the experimental setup: SCC25 human squamous cell carcinoma cells were treated with the pan-Notch inhibitor CB103 or DMSO (control) for five days prior to downstream analyses. (B) qRT-PCR analysis shows that CB103 treatment significantly decreases expression of the Notch ligand DLL4 and the stemness-associated markers SOX2 and P63. (C) Bar plot summarizing Gene Ontology (GO) classification of cellular components, comparing gene expression profiles between CB103- and DMSO-treated SCC25 cells. (D) Heatmap with hierarchical clustering of the top 500 differentially expressed genes identified via bulk RNA sequencing of SCC25 cells treated with CB103 or DMSO. (E) Gene Set Enrichment Analysis (GSEA) plots show enrichment of keratinization (left) and keratinocyte differentiation (right) signatures following CB103 treatment. (F) Volcano plot highlighting upregulation of keratin genes (e.g., KRT1, KRT10) and downregulation of the stemness marker SOX2 after CB103 treatment. (G) GSEA plot showing increased expression of genes involved in keratin filament organization in CB103-treated cells. (H) GSEA plot illustrating changes in genes associated with cytoskeletal components following CB103 exposure. (I) GSEA plot showing modulation of the epithelial-to-mesenchymal transition (EMT) gene signature in response to CB103 treatment.

To further investigate the molecular impact of CB103 treatment, we performed bulk seqRNA comparing DMSO- versus CB103-treated hSCC25 cells. This analysis identified 3329 differentially expressed genes (p≤ 0.01; Supplementary Method 2), with the top 2000 genes visualized in a hierarchical clustering heatmap (Figure 8D). The heatmap highlights alterations in genes involved in epithelial differentiation (Cluster 3, 4 and Cluster 5), cell-to-cell adhesion (Cluster 3), cytoskeleton organization (Cluster 4, Cluster 5) and chemotaxis (Cluster 1) (Figure 8D, Supplementary Table 4). GO analysis for cellular components and biological processes, along with *Enrichr* analyses, further supported these findings by identifying alterations in intermediate filaments, cell-to-cell adhesion, and cell-to-ECM interactions (Figure 8C, Supplementary Figure 7A and Supplementary Figure 7B). Paralleling our findings in the mouse model, GSEA analysis revealed significant changes in keratinization processes and keratinocytes differentiation (Figure 8E). Additionally, MA and volcano plot analysis indicated an upregulation in the expression of keratins (Figure 8F), as well as a change in the expression of integrins and ECM molecules (Supplementary Figure 7C). Together with GSEA results (Figure 8G-I), these data suggest that CB103-treated cells undergo structural modifications and remodeling of anchoring molecules.

We hypothesized that such changes could affect cell motility, a crucial factor in acquiring an invasive phenotype during cancer progression. We therefore aimed to explore the functional consequences of CB103 treatment. Among the differentially expressed signatures, we identified enrichment in genes associated with desmosomal complexes and apical junctions (Figures 9A, 9B and 9D). qRT-PCR validation confirmed a significant upregulation of desmosome-associated proteins upon CB103 treatment, including desmoplakin, desmoglein, and desmocollin (Figure 9C). Consistently, CB103-treated cells exhibited a marked increase in the number of adhesive complexes compared to DMSO-treated controls (Figures 9E and 9F)^68,69^, reinforcing the role of Notch inhibition in enhancing cell–cell adhesion. Finally, we conducted scratch assays and monitored the progression of migrating cells over a period of 10 days (Figures 9G and 9H). DMSO-treated cells were able to fully reform the monolayer within a ten-day timeframe, in contrast to CB103-treated cells, which were unable to recolonize the cell-free zone (Figure 9G). To further confirm the reduced motility of CB103-treated cells, we performed time-lapse analysis. CB103-treated cells exhibited significantly reduced motility when compared to cells treated with DMSO (Figures 9I and 9J; Movies 4 and 5). These results indicate that selective Notch inhibition by CB103 markedly alters the expression of structural components within adhesive complexes that anchor epithelial cells to the extracellular matrix, ultimately reducing the migration of human carcinoma cells.

**Figure 9:**
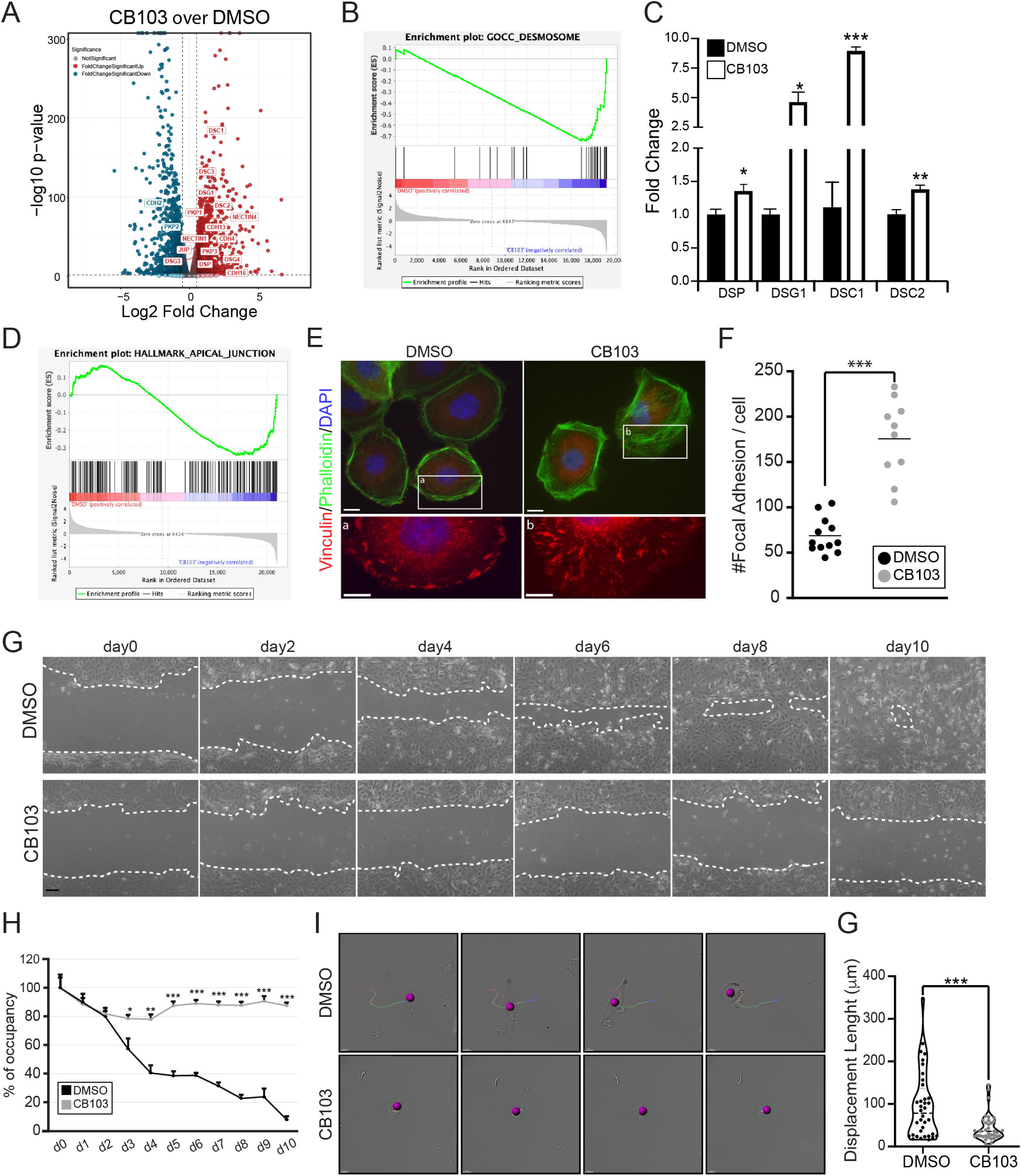
Blockage of Notch transcription results in disruption of SCC cells migratory activity. A) Volcano plot showing upregulation of transcripts associated with desmosomes and cell adhesion molecules in CB103-treated cells (e.g., plakoglobin (JUP), desmocollins (DSC1/2), and desmogleins (DSG1/2). (B) GSEA plot highlighting enrichment of gene ontology cellular components related to desmosomes in CB103- versus DMSO-treated conditions. (C) qRT-PCR analysis of SCC25 cells treated with DMSO or CB103 demonstrates increased expression of desmosomal structural components following Notch inhibition (e.g. desmoplakin (DSP), desmoglein 1 (DSG1), and desmocollins (DSC1 and DSC2)). (D) GSEA plot showing changes in apical junction-related gene signatures in SCC25 cells treated with CB103. (E) Representative immunofluorescence images of actin filaments and focal adhesions in SCC25 cells treated with DMSO or CB103. Green: phalloidin. Red: vinculin. Blue: DAPI. Scale bar: 10 µm. (F) Quantification of focal adhesion complexes per cell. *** p < 0.001. (G) Scratch wound assay of SCC25 cells treated with DMSO or CB103. Scale bar 100μm. (H) Quantification of the cell-free area in the scratch assay. Values normalized to the initial scratched area (day 0). Error bars represent standard error. **=p<0.01; ***=p<0.001; ****=p<0.0001 (I) Time-lapse imaging of SCC25 cells treated with DMSO or CB103, showing single frames from representative recordings over a 2-hour 25-minute period (related to Movies 3 and 4). Scale bar 50μm. (J) Quantification of single-cell displacement lengths using Imaris spot tracking.***=p<0.001

Overall, our findings from both murine and human models highlight the critical role of the Notch1-Dll4 axis in maintaining undifferentiation and promoting motility. Disrupting this pathway effectively impairs key drivers of OSCC malignancy.

## Discussion

The Notch signaling pathway is a central regulator of carcinogenic transformation in a variety of epithelial tumors, including lung, cervical, colon, and pancreatic cancers ^70–73^. The dysregulation of Notch is context-dependent and can shift from an activator to an inhibitor, depending on the organ and the type of tissue affected ^74–76^. Currently, a debate exists on whether the Notch pathway acts as an oncogene or tumor suppressor in OSCC, and its exact role in this type of cancer remains largely unknown ^19,77,78^. Since Notch activity is strongly influenced by the specific receptor-ligand interactions within a tissue, our first goal was to identify how these combinations evolve across healthy tissue, initiation, and advanced stages of OSCC. To achieve this, we used the *Notch1CreERT:R26mTmGfl/fl* transgenic line and studied Notch1-expressing cells and their progeny throughout the progressive stages of malignant transformation. Short tamoxifen pulses allowed precise control of the reporter activation, providing information on which cells had active Notch1 at specific time points. Pulse-and-chase experiments instead, tracked labelled cells over extended periods, revealing their movements and dynamic behavior as the cancer progresses and its structure evolves. Through detailed phenotypic characterization, our findings suggest that Notch signaling contributes to OSCC development and progression by preferentially interacting with the Dll4 ligand rather than Jag1. Gene expression analyses confirmed that *Dll4* is the most significant gene of the Notch pathway to be differentially regulated in OSCC condition. The preferential binding of Notch1 to Dll4 in the tumorigenic epithelium ultimately leads to the downregulation of *Hey1* and *Hey2*, while the expression of other Notch-specific target genes, such as *Hes1* and *Hes2,* remains largely unaffected. The mutually exclusive expression of the Jag1 and Dll4 ligands regulates fate definition and undifferentiation in a variety of tissues (e.g., during angiogenesis and hematopoiesis)^40,79,80^. Similarly in our system, the selection between the Notch1-Dll4 versus Notch1-Jag1 interaction observed in OSCC coincides with sustained keratinocyte undifferentiation. The Notch1/Dll4-expressing keratinocyte progenitors retain their immature state, resulting in the expansion of the undifferentiated portion of the tongue epithelium.

Important downstream changes occur during Notch pathway dysregulation in OSCC. Our transcriptomic analyses reveal a complex scenario of crosstalk between pathways sustaining oral carcinoma. Interestingly, p63 and Runx1 cooperate with the Notch pathway, similarly to what has been found in other tissues. In skin keratinocytes a bidirectional crosstalk between Notch and p63 suggest a complex regulation of differentiation occurring at various levels (from regulation of transcription to indirect regulation of the Notch pathway). A high expression of P63 correlates with poor prognosis in patients with OSCC of the tongue^53,54,64,81–83,75^. Our model showed a similar correlation, with an undifferentiated mass coherently expressing P63 and Notch1-Dll4. In HNSCC screenings, the transcription factor RUNX1 was found to be significantly increased and thus, has been suggested as a prognostic marker ^83,84^. RUNX1 upregulation drives tumor progression and metastasis, while suppression of RUNX1 inhibits HNSCC cell proliferation and migration ^84,84,85^. In other settings, such as mammalian hematopoiesis, Runx1 controls the expansion, specification and self-renewal of stem cells in a Notch-dependent manner ^86^ ^87,88^. We have found a similar scenario in our system, where both transcriptomic approaches and immunostainings show a widespread expression of Runx1 and Notch1-Dll4 with an analogous expression pattern.

Continuous epithelial growth gives rise to a malignant mass with the potential of invading the underling stroma^11^. In the invasive growth, we identified typical features of EMT, such as basal lamina dissolution, overexpression of ZEB1, ZEB2 and Snail1 and cells co-expressing Vimentin and E-cadherin. Transcriptomic analysis confirmed a specific hallmark signature for EMT in the epithelium of 4NQO-treated mice. Importantly, cells of epithelial origin growing within the mesenchyme maintain the expression of Dll4. Consequently, we identified the Notch-Dll4 axis throughout all phases of OSCC progression, spanning from the initial hyperplastic phase to the most advanced invasive stages, highlighting its promising potential as a diagnostic tool.

To strengthen the translational impact of our findings, we analyzed human malignant tissue and confirmed elevated expression of DLL4. Dll4 blockers have been proposed for the treatment of other epithelial tumors (e.g., intestinal tumors) primarily aiming to limit angiogenesis ^89^. Since OSCC growth is not strictly dependent on vascularization, a locally administered blocker could directly target malignant epithelial cells in this tumor. Immunotherapy specifically targeting Dll4 combined to chemotherapy has been applied to treat solid tumors (e.g., pancreatic and lung cancers) ^90–93^, with positive results spanning from a reduction of tumor growth to a decrease in the reoccurrence rate ^94–96^. Given the redundancy of the Notch pathway and the frequent activation of compensatory mechanisms when a single component is targeted, we hereby adopted a broader strategy to test the therapeutic potential of pan-Notch blockers in OSCC. The Notch pharmacological inhibitor CB103 is increasingly gathering attention in cancer treatments, in view of its high stability and unbeaten specificity in targeting the Notch pathway ^67,97,98^. Several clinical trials are currently in place to determine its usage as a therapeutic agent in carcinomas (clinical trials: NCT03422679, NCT04714619) as well as leukemias and lymphomas (clinical trials: NCT03422679, NCT05464836). In our system, SCC25 cells treated with CB103 displayed a significant reduction in *DLL4* expression, with no observable change in *JAG1* expression, confirming the crucial role of the NOTCH-DLL4 signaling axis in OSCC. Importantly, the expression of undifferentiation markers SOX2 and P63 is significantly reduced in CB103-treated cells, while transcriptomic analysis reveals induction of keratinization. These *in vitro* results validate the *in vivo* findings in mice and denote the connection between an active NOTCH-Dll4 signaling axis and the prolonged undifferentiation cell state within OSCC. We finally tested the functional effects of Notch blockage in OSCC cell behavior. We recorded a dramatic reduction in cell migration when SCC25 cells were treated with CB103. This is accompanied by an increase in adhesive complexes (focal adhesions and desmosomes), suggesting that NOTCH regulates cell movement and is essential for the acquisition of tumor’s invasive properties.

In conclusion, the present findings show that the Notch1-Dll4 signaling axis plays a crucial role in the initiation and development of OSCC, contributing to the acquisition of critical oncogenic features.

## Supporting information

Sup Info_Meisel et al.

Sup Figure 1_Meisel et al.

Sup Figure 2_Meisel et al.

Sup Figure 3_Meisel et al.

Sup Figure 4_Meisel et al.

Sup Figure 5_Meisel et al

Sup Figure 6_Meisel et al.

Sup Figure 7_Meisel et al

Movie 1_meisel et al

Movie 2_Meisel et al

Movie 3_meisel et al

Movie 4_Meisel et al

Movie 5_Meisel et al

## Acknowledgements

This work was supported by the University of Zurich, the Krebsliga (KFS-4890-08-2019-R) (T.A.M.), the Julius Klaus-Stiftung (C.P.), Stiftung Julius Müller zur Unterstützung der Krebsforschung (C.P.) and the Olga Mayenfisch Stiftung (C.P.). We thank Prof. Spyros Artavanis-Tsakonas (Harvard Medical School, Harvard University) for providing *Notch1^creER^*mice.

Imaging was performed with equipment maintained by the Center for Microscopy and Image Analysis, University of Zurich. The authors gratefully acknowledge the Functional Genomics Center Zurich (FGCZ) of University of Zurich and ETH Zurich for the support on analyses.

## List of abbreviations

4NQO: 4-nitroquinoline-1-oxide
CSL: CBF, Suppressor of Hairless, Lag-1
Dll4: Delta-like-4
DMSO: Dimethyl sulfoxide
E-Cad: E-cadherin
EMT: Epithelial-to-mesenchymal-transition
GFP: Green fluorescent protein
GO: Gene Ontology
GSEA: Gene set enrichment analysis
Hes: Hairy and enhancer of split-1
Hey: Hairy and enhancer of split related with YPRW motif protein
HNSCC: Head and neck squamous cell carcinoma
Jag1: Jagged1
K5/10/14: Keratin 5/10/14
MesoSPIM: Mesoscale selective plane illumination microscopy
OSCC: Oral squamous cell carcinoma
RBPj: Recombining binding protein suppressor of hairless
Runx1: Runt-related transcription factor 1
Sox2: SRY-Box Transcription Factor 2
TAM: Tamoxifen

