## Supplementary material for "The Notch1/Delta-like-4 axis is crucial for the initiation and progression of oral squamous cell carcinoma": Sup Info_Meisel et al.

**Supplementary Figure 1: Chronic exposure to 4NQO reproduces in mice the human progressive evolution of OSCC**

(A) Schematic representation of mice treated with DMSO or 4NQO. 6 weeks-old mice received drinking water containing DMSO (control group) or 4NQO (treated group) continuously for 16 weeks followed by normal water. Mice were euthanized at 22 weeks of age (early time point) or 32 weeks of age (advanced time point) for dissection and processing of the tongue organ. (B) Brightfield images of whole tongues post-treatment from 22 weeks-old mice (left panels) and 32 weeks-old mice (right panels). (C) Histological analysis of the dorsal part of tongues treated with DMSO/4NQO from 22 weeks-old mice (upper panels) and 32 weeks-old mice (lower panels). (*)

Indicates magnified area in the corresponding region of the overview. Scale bars in overviews 100 µm.□Scale bars in magnification 50µm.

**Supplementary Figure 2: The 4NQO overgrowth is formed by undifferentiated progenitor cells**

(A) Cross-section stained against Sox2 (green) and DAPI (blue), displaying the dorsal part of the tongue after DMSO treatment (left panels) and after 4NQO treatment (right panels). (*) and (**) Indicate magnified areas in the corresponding regions of the overview. (B) Cross-section stained against K5 (red) and DAPI (blue), displaying the dorsal part of the tongue after DMSO treatment (left panels) and after 4NQO treatment (right panels). (*) and (**) Indicate magnified areas in the corresponding regions of the overview. (C) Cross-section stained against K10 (cyan) and DAPI (blue), displaying the dorsal part of the tongue after DMSO treatment (left panels) and after 4NQO treatment (right panels). (*) and (**) Indicate magnified areas in the corresponding regions of the overview. (D) Cross-section stained against E-Cadherin (green) and DAPI (blue), displaying the dorsal part of the tongue after DMSO treatment (left panels) and after 4NQO treatment (right panels). (*) and (**) Indicate magnified areas in the corresponding regions of the overview. Scale bars in overviews100µm. Scale bars in magnification 50µm.

**Supplementary Figure 3: GSEA identifies changes in the gene signature for differentiation**

(A) Gene Set Enrichment Analysis (GSEA) identifies multiple transcriptomic signatures for changes in keratinocytes differentiation, keratinization and specific taste-bud fate acquisition.

**Supplementary Figure 4: Transcriptome analysis of downstream pathway**

(A) Connection plot between differentially expressed groups of genes. *Runx1* regulation is highlighted in red. (B) Box-plot diagram for genes of the *Runx*-family and *p21*. *Runx1* changes of expression is the most prominent amongst the Runx genes. (B) *Enricher* analysis of pathways identifies *Ep300*, *Stat3* and *Runx1* in between the most altered genes. (D) *Enricher* independent analyses of transcription factor interaction confirms alterations of STAT3 and ep300.

**Supplementary Figure 5: Regulation of Runx1 and P63 in 4NQO-treated tissue**

(A) Cross-sections displaying the dorsal part of the tongue stained against Runx1 (green) after DMSO- (left panels) or 4NQO- (right panels) treatment. Nuclear counterstaining with DAPI (Blue). (*) Indicates magnified area in the corresponding region of the overview. (B) Sections displaying the dorsal part of the tongue stained against P63 (green) after DMSO- (left panels) or 4NQO- (right panels) treatment. (*) and (**) Indicate magnified areas in the corresponding regions of the overview. Nuclear counterstaining with DAPI (Blue). Scale bars in overviews 100µm. Scale bars in magnification 50µm.

**Supplementary Figure 6: Light-sheet microscopy illustrates the early phases of basal lamina dissolution.** (A) SPIM analysis of the entire thickness of the tongue organ in 22-week-old 4NQO-treated mice. The basal lamina preserves its integrity throughout the digital sectioning (yellow arrows). (B) SPIM analysis of the entire thickness of the tongue organ in 32-week-old 4NQO-treated mice. Digital sectioning highlights a distinct area with disrupted basal lamina (white arrows). Scale bar 200 µm.

**Supplementary Figure 7:Transcriptome analysis of SCC25 cells treated with CB103 Notch blocker**

(A) Gene Ontology (GO) classification analysis for categories of biological processes comparing CB103 over DMSO controls. Note the main changes in cell adhesion, keratinization and differentiation. (B) Clustergram from Enrichr analyses GOCC. Cornified envelope, keratin filaments and desmosomes genes are the top enriched terms. (C) MA plots for integrins (upper panel) and extracellular-matrix components (lower panel) show log-fold change of genes against average expression.

**Supplementary Movie 1: Three-dimensional expression of Notch1-driven GFP in the murine tongue epithelium**

Movie showing the GFP channel (green) of a *Notch1*$^{CreERT}$*:R26mTmG*$^{fl/fl}$ mouse tongue upon Tamoxifen-induced Cre-activation. The rotation on central axis demonstrates the expression of Notch1-driven GFP in dorsal and ventral epithelium of the tongue. Scale bar 300µm

**Supplementary Movie 2: Whole-organ imaging to study basal lamina integrity**

The movie shows the background fluorescence of a 22-week-old 4NQO-treated murine tongue. The section viewer enables visualization of digital sections along the Y-axis to analyze the entire thickness of the organ. At this stage we identified a growing malignant mass lying on an intact basal lamina. Related to Supplementary Figure 3. Scale bar 200µm.

**Supplementary Movie 3: Whole organ imaging reveal dissolution of basal lamina**

The movie shows the background fluorescence of a 32-week-old 4NQO-treated murine tongue. The section viewer enables visualization of digital sections along the Y-axis to analyze the entire thickness of the organ. We identified a specific region where the basal lamina is degraded, indicating a carcinoma with invasive potential. Related to Supplementary Figure 3. Scale bar 500µm.

**Supplementary Movie 4: Time-lapse analysis of DMSO-treated SCC25 cells.**

Brightfield transmission recording over 2h25' with an image capture every 3'. DMSO-treated SCC25 cells are tracked via Spot&Tracking function in Imaris 9.9 software (colored sphere as central point) for the calculation of the displacement length. Tracking over-time progression visible via the rainbow-colored trail (from blue to red). Representative snapshots in Fig 7G. Scale bar 50µm.

**Supplementary Movie 5: Time-lapse analysis of CB103-treated SCC25 cells.**

Brightfield transmission recording over 2h25' with an image capture every 3'. CB103-treated SCC25 cells are tracked via Spot&Tracking function in Imaris 9.9 software (colored sphere as central point) for the calculation of the displacement length. Tracking over-time progression visible via the rainbow-colored trail (from blue to red). Representative snapshots in Fig 7G. Scale bar 50µm.

**Supplementary Method 1: Threshold of significance settings for transcriptome analyses on murine OSCC model**

(A) General settings for transcriptomic RNA sequencing. (B)Threshold significance was set to $p<0.01$ corresponding to FDR level of 0.045. (C) Significant changes in expression were found for 3331 genes. (D) Volcano plot and average expression of transcripts. Significant genes were depicted as blue circles.

**Supplementary Method 2: Threshold of significance settings for transcriptome analyses on human SCC25 cells**

(A) General settings for transcriptomic RNA sequencing. (B)Threshold significance was set to $p<0.01$ corresponding to FDR level of 0.017. (C) Significant changes in expression were found for 3329 genes. (D) Volcano plot and average expression of transcripts. Significant genes were depicted as blue circles.

**Supplementary Table 1: Murine primer pairs for quantitative real-time PCR**

| **Target** | **FW primer 5’-3’** | **RV primer 3’-5’** |
|---|---|---|
| mActin | CAT TGC TGA CAG GAT GCA GAA GG | TGC TGG AAG GTG GAC AGT GAG G |
| mGapdh | TGCGACTTCAACAGCAACTC | ATGTAGGCCATGAGGTCCAC |
| mJag1 | TTCTCACTCAGGCATGATAAACC | CATCTCTGGGACGACAGAACT |
| mDll4 | GGAACCTTCTCACTCAACATCC | CTCGTCTGTTCGCCAAATCT |
| mHes1 | GGAAATGACTGTGAAGCACCTCC | GAAGCGGGTCACCTCGTTCATG |
| mHes2 | TCAACGAGAGCCTAAGCCAGCT | CGCACAGTCATTTCCAGGATGTC |
| mHey1 | CCAACGACATCGTCCCAGGTTT | CTGCTTCTCAAAGGCACTGGGT |
| mHey2 | TGAAGATGCTCCAGGCTACAGG | CCTTCCACTGAGCTTAGGTACC |

| | | |
|---|---|---|
| mHeyL | CTGGAGAAAGCTGAGGTCTTGC | ACCTCAGTGAGGCATTCCCGAA |

**Supplementary Table 2: Human primer pairs for quantitative real-time PCR**

| **Target** | **FW primer 5’-3’** | **RV primer 3’-5’** |
|---|---|---|
| hActin | CAC CAT TGG CAA TGA GCG GTT C | AGG TCT TTG CGG ATG TCC ACG T |
| hGAPDH | AGCCACATCGCTCAGACAC | GCCCAATACGACCAAATCC |
| hJag1 | TGCTACAACCGTGCCAGTGACT | TCAGGTGTGTCGTTGGAAGCCA |
| hDll4 | CTGCGAGAAGAAAGTGGACAGG | ACAGTCGCTGACGTGGAGTTCA |
| hP63 | CAGGAAGACAGAGTGTGCTGGT | AATTGGACGGCGGTTCATCCCT |
| hDSP | GCCCTGTGATGCTTACCAGA | GCCTCATCCACCCCAAACAT |
| hDSG1 | AGCTCTGAACTCAATGGGCC | TGTTCGGTTCATCTGCGTCA |
| hDSC1 | GCAAGAGACGATGGGCTCCT | AGGGTTCTTTGTCCACGCCT |
| hDSC2 | CCTGGATAGAGAGGCAGAGACC | CAACCGCAACAATCTCCGCA |
