## Supplementary material for "The Notch1/Delta-like-4 axis is crucial for the initiation and progression of oral squamous cell carcinoma": Sup Figure 1_Meisel et al.

Supplementary Figure 1: Chronic exposure to 4NQO reproduces in mice the human progressive evolution of OSCC

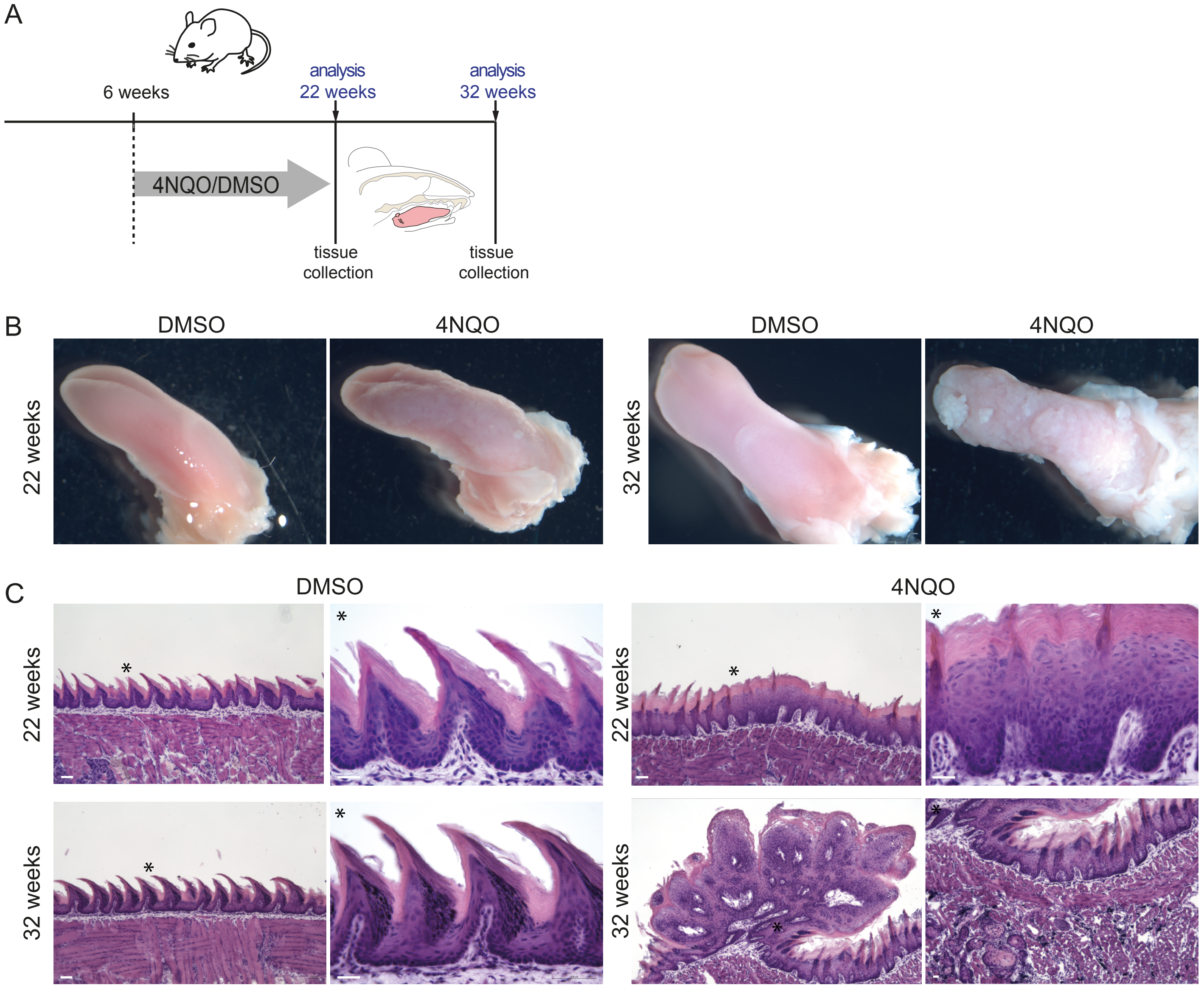
