## Supplementary figures and images for "The Notch1/Delta-like-4 axis is crucial for the initiation and progression of oral squamous cell carcinoma"

### Sup Figure 2_Meisel et al.

Supplementary Figure 2: The 4NQO overgrowth is formed by undifferentiated progenitor cells

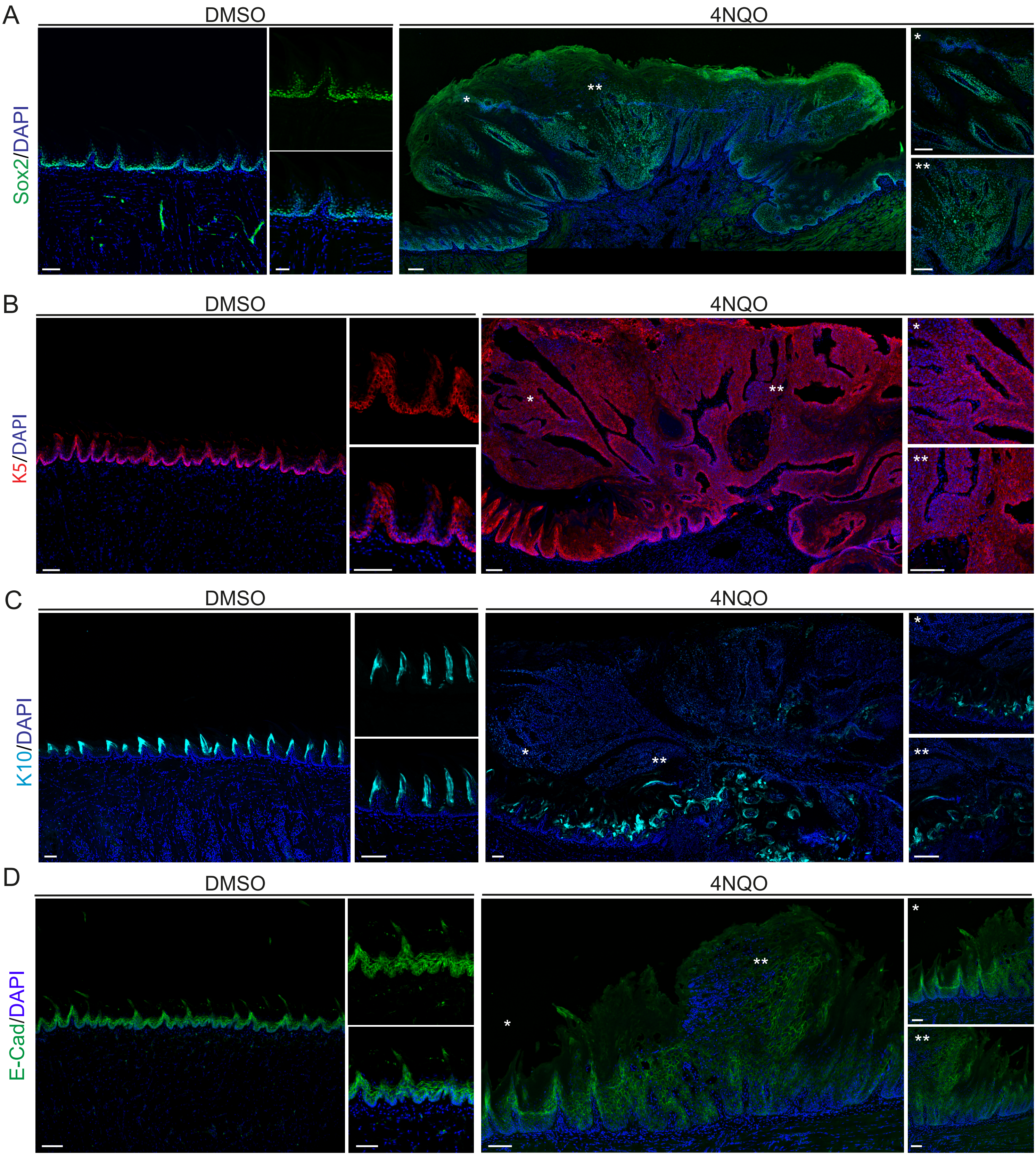

### Sup Figure 3_Meisel et al.

Supplementary Figure 3: GSEA identifies changes in the gene signature for differentiation

A

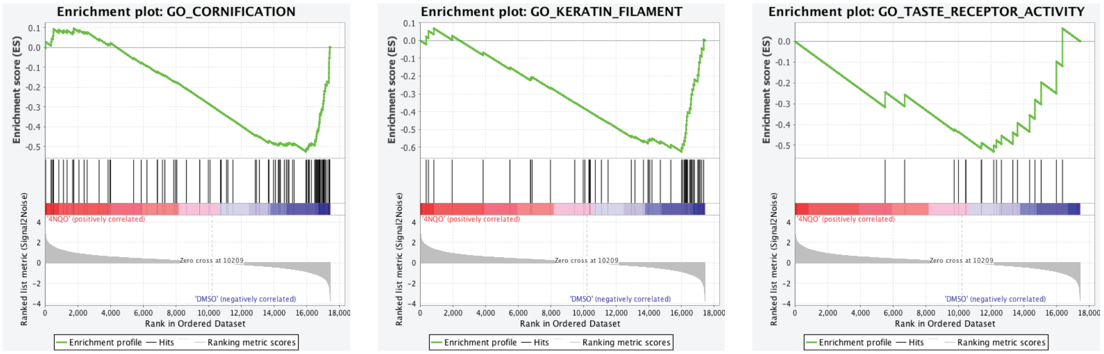

### Sup Figure 5_Meisel et al

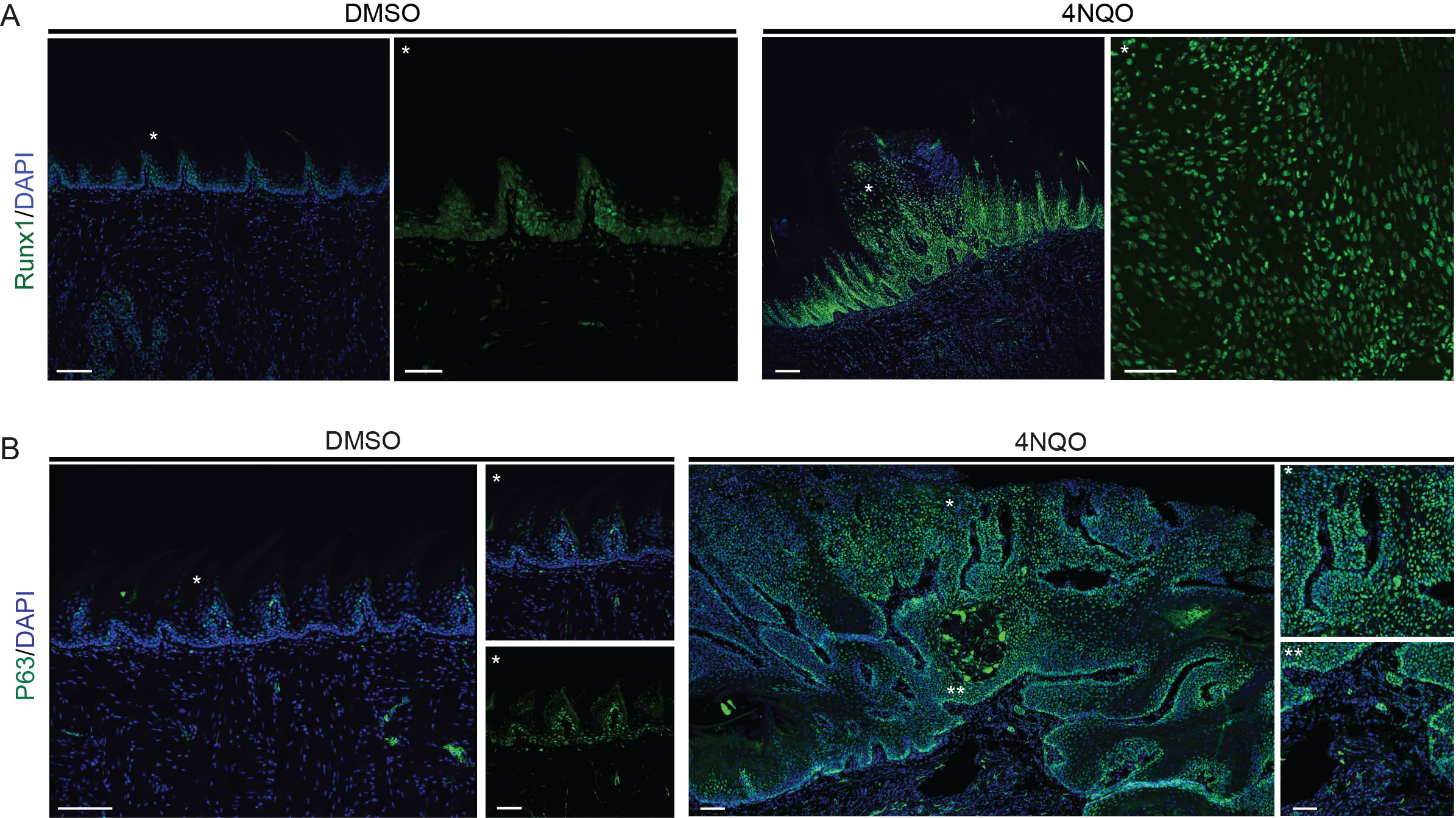

### Sup Figure 7_Meisel et al

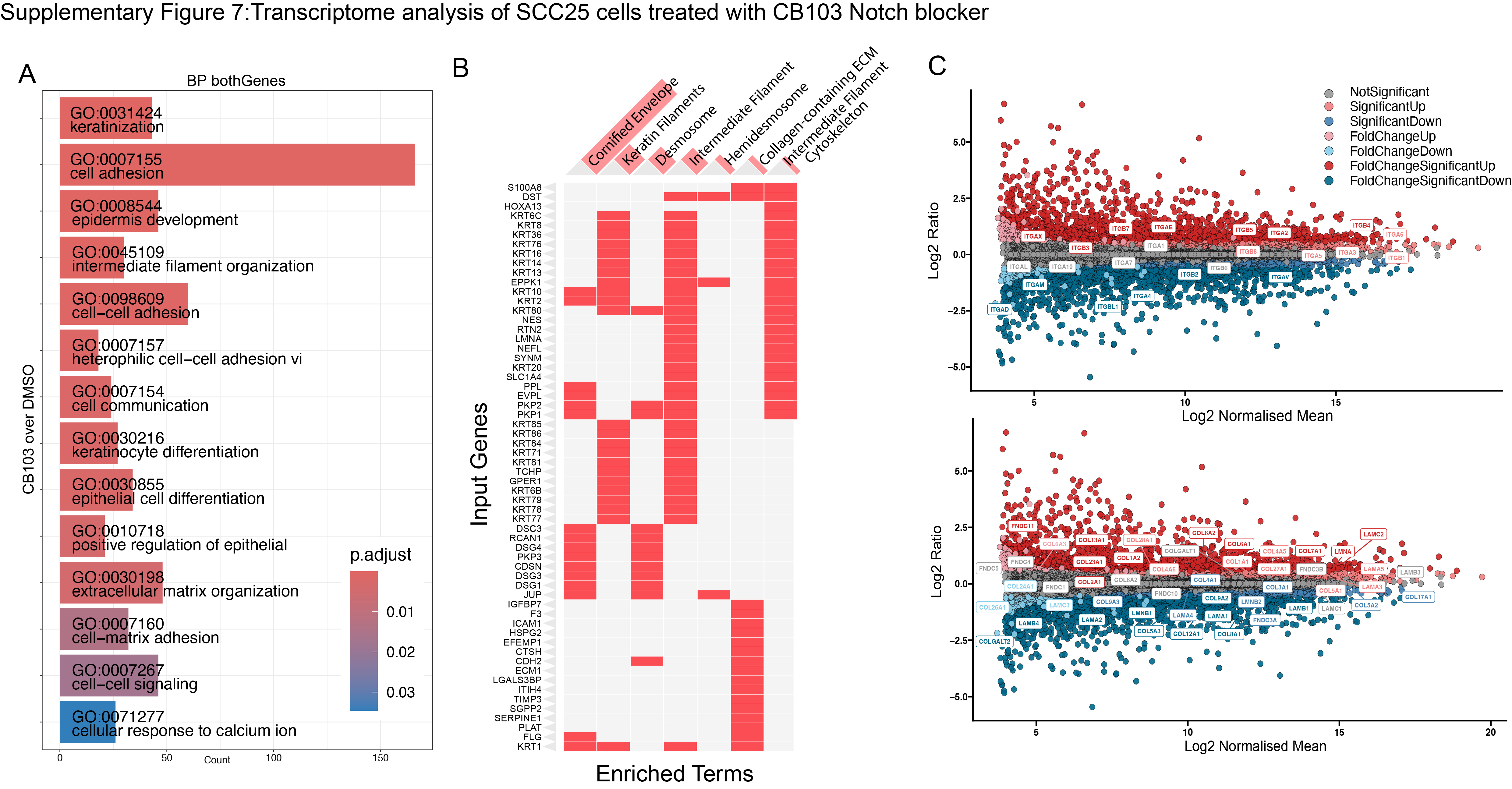
