## Supplementary material for "The Notch1/Delta-like-4 axis is crucial for the initiation and progression of oral squamous cell carcinoma": Sup Figure 4_Meisel et al.

### Supplementary Figure 4: Transcriptome analysis of downstream pathway

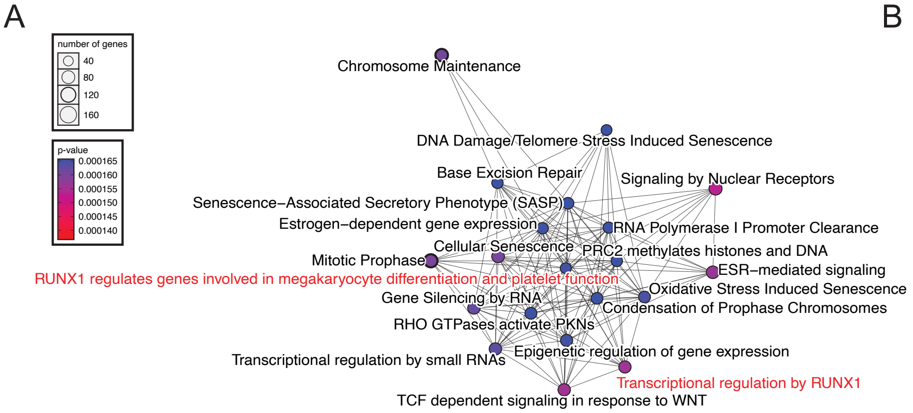


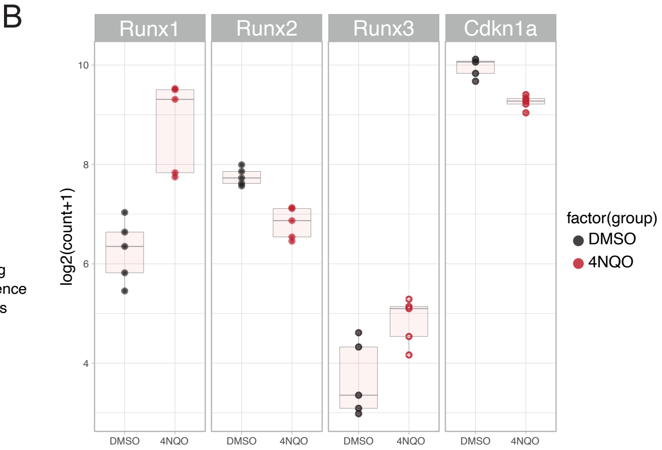


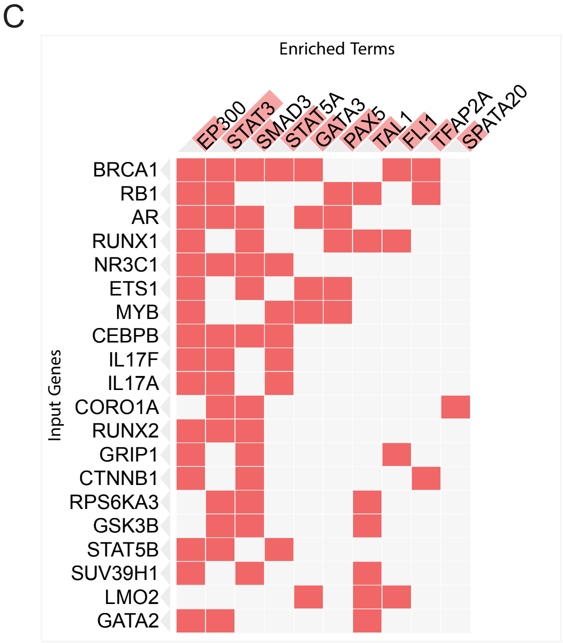


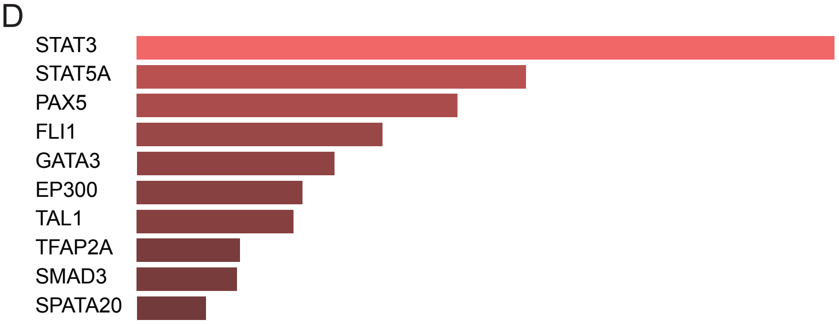


| Index | Name | P-value | Adjusted p-value | Odds Ratio | Combined score |
|---|---|---|---|---|---|
| 1 | STAT3 | 0.00008910 | 0.02317 | 1.78 | 16.59 |
| 2 | STAT5A | 0.001564 | 0.2034 | 2.05 | 13.23 |
| 3 | PAX5 | 0.002958 | 0.2563 | 4.72 | 27.49 |
| 4 | FLI1 | 0.005922 | 0.3849 | 3.54 | 18.16 |
| 5 | GATA3 | 0.009290 | 0.4831 | 2.95 | 13.81 |
| 6 | EP300 | 0.01248 | 0.5036 | 1.30 | 5.72 |
| 7 | TAL1 | 0.01356 | 0.5036 | 1.95 | 8.39 |
| 8 | TFAP2A | 0.02231 | 0.6608 | 2.07 | 7.86 |
| 9 | SMAD3 | 0.02287 | 0.6608 | 1.31 | 4.93 |
| 10 | SPATA20 | 0.03064 | 0.7472 | 32998.00 | 115018.12 |
