## Supplementary material for "The Notch1/Delta-like-4 axis is crucial for the initiation and progression of oral squamous cell carcinoma": Sup Figure 6_Meisel et al.

Supplementary Figure 6: Light-sheet microscopy illustrates the early phases of basal lamina dissolution.

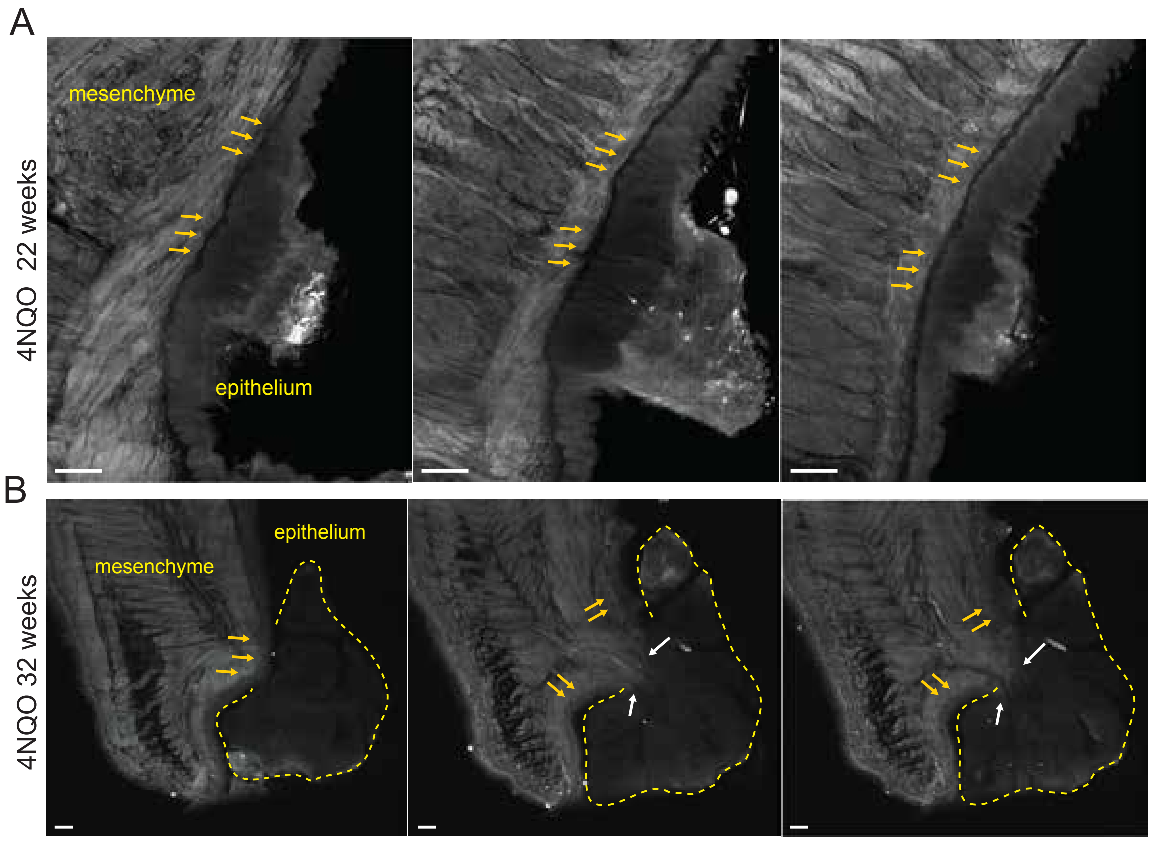
